# Sustained MYB activity drives emergent enhancer activation and precise enhancer-promoter interactions

**DOI:** 10.1101/2025.04.16.649256

**Authors:** I-Jun Lau, Joe R. Harman, Alastair L. Smith, Nicholas Denny, Nicole E. Jackson, Joseph C. Hamley, Paresh Vyas, James O.J. Davies, Jim R. Hughes, Nicholas T. Crump, Thomas A. Milne

**Author notes:** **Correspondence:** (T.A.M.), (N.T.C.).

## Abstract

Transcription factors (TFs) are key effectors of enhancer activity. MYB is a critical hematopoietic TF that is frequently dysregulated in cancer. Despite its well-established role, the exact mechanisms by which MYB influences enhancer function—and the specific stages of enhancer activation at which it operates—remain poorly understood. Using high resolution Micro-Capture-C, we show that upon MYB degradation, highly defined enhancer-promoter interactions at specific MYB binding sites are lost. Loss of these interactions, together with other hallmarks of enhancer activity—reduced H3 lysine-27 acetylation and enhancer RNA transcription—correlates with significant downregulation of target gene expression in leukemia, indicating that MYB mediates transcription activation via maintenance of enhancer function. When anchored to DNA within a gene desert region that is devoid of histone marks and active transcription, the MYB transactivation domain is sufficient and necessary for the nucleation of an enhancer-like region. This results in the activation of transcription from distal cryptic elements and the establishment of long-range chromatin interactions up to 400 kb away from the anchor point. Together, these results indicate that MYB activity alone is sufficient to induce long-range interactions and transcription, achieving this through highly precise enhancer-promoter crosstalk.

## Introduction

The precise regulation of gene expression is essential for life. Disruption of normal gene transcription is a major driver of human disease, including cancer. Transcription factors (TFs) are a class of proteins that are responsible for the complex spatiotemporal and tissue-specific control of gene expression (*1, 2*). TFs bind to promoter-distal cis-regulatory elements known as enhancers and once bound, modulate expression of their target genes from these distal loci, thereby facilitating the orchestration of differential gene expression programs (*3–5*). The clustering of TF binding is a key feature of active enhancers, alongside histone modifications such as H3 lysine-27 acetylation (H3K27ac) and H3 lysine-4 monomethylation (H3K4me1), as well as bidirectional non-coding, enhancer RNA (eRNA) transcription (*5–8*). Although there are examples where co-localization of the enhancer and promoter does not appear to be important for gene activation (*9, 10*), in most instances enhancer-promoter proximity is a key measure of enhancer function (*11–17*), even if it is unclear what drives these interactions. Co-activators such as Mediator, BRD4 and P300/CBP have been implicated in enhancer activity and are believed to cooperate with TFs in order to induce enhancer activation (*2, 18, 19*). However, mounting evidence suggests that many co-activators have only subtle impacts on enhancer-promoter interactions (*20–22*) and H3K27ac is dispensible for enhancer activity in embryonic stem cells (ESCs) (*7*). Current high-resolution 3C methods such as Region Capture Micro-C (RCMC) or Micro-Capture-C (MCC) have made the striking observation that enhancer-promoter interactions are precisely localized to TF binding sites, raising the possibility that enhancer-promoter crosstalk is driven by TF interactions (*23–25*). Recently, the transcription cofactor LDB1 has been shown to be essential for loop formation (*17*), but it is unclear if this mainly drives enhancer-promoter crosstalk or impacts higher order chromatin structure, or how universal this activity may be among key TFs (*26*). Thus, despite the importance of enhancers and the TFs that bind to them, how TF effector function and enhancer activity are coupled, and which aspects of enhancer activity are TF-dependent remain unknown.

MYB is a developmentally important TF that is required for definitive hematopoiesis (*27–29*). One of the main activities of MYB is to recruit the lysine acetyltransferases P300/CBP to enhancers, increasing localized H3K27ac levels. Mutations within the transactivation domain of MYB that disrupt its interaction with the KIX domains of P300/CBP (*30–33*) and vice versa, lead to significant hematopoietic defects and prevent the induction of leukemia by known oncogenes in mouse models. In addition to leukemia (*34*), aberrant MYB activity has been implicated in a wide range of cancers, as well as contributing to drug resistance (*35*). Spontaneous formation of a new binding site for MYB is sufficient to create a novel, active enhancer that drives leukemia (*36*), at least in part by activating cryptic binding sites for other TFs in the same region, which then act as cooperative partners with MYB. However it is not clear whether MYB alone is able to create a new regulatory unit.

To address this gap in knowledge, we investigated the relationship between enhancer activity and MYB binding by introducing a degradation tag (FKBP12^F36V^) to the endogenous *MYB* gene to allow rapid protein loss in leukemia cells. By coupling this with the high resolution 3C technique MCC, we show that overall enhancer-promoter looping is maintained upon MYB degradation, but highly defined interactions at MYB binding sites are lost. These interaction losses were associated with significant transcriptional downregulation of known MYB target genes and correlated with decreased enhancer H3K27ac levels and eRNA transcription. To interrogate MYB function and determine whether it can recruit specific enhancer activities de novo, we anchored the Myb transactivation domain (Myb^TA^) to a gene desert region and found it was sufficient to recruit key enhancer-associated proteins, including P300, Brd4 and Mediator to chromatin. We observed Myb^TA^-dependent deposition of H3K27ac, increased chromatin accessibility and transcription – not just at the anchor site, but remarkably, also at regions more than 50 kb away, likely cryptic promoters. Myb^TA^ induced DNA looping to facilitate 3D interactions with these distal sites of transcription, as well as more long-range interactions spanning up to 400 kb.

Overall, our data argue for a role for MYB in establishing enhancers de novo, by directing specific protein-protein interactions from distal regulatory elements. At a subset of MYB-bound enhancers, the continued presence of MYB is absolutely required to maintain enhancer activity and oncogene upregulation. These observations not only provide a specific model for MYB function, but also provide a paradigm for understanding the general function of enhancers.

## Results

### MYB binds to active enhancers in leukemia cells

To assess the role of MYB in enhancer function, we performed a genome-wide analysis in the B leukemia cell line SEM. MYB binding positively correlated with enhancer-associated factors P300, BRD4 and MED1, as well as RNAPII and H3K27ac, both globally (Fig. S1A) and specifically at enhancers (Fig. 1A, B). We used Next Generation Capture-C (*37*) to visualize 3D interactions with the promoters of key MYB-dependent oncogenes including *BCL2*, *MYC, LMO4* and *FLT3*, identifying both intergenic and intragenic enhancers bound by MYB (Fig. 1C, D, S1B, C). With comparatively low levels at the promoters of these genes, this suggests an enhancer-centered role for MYB in their expression.

**Fig. 1.**
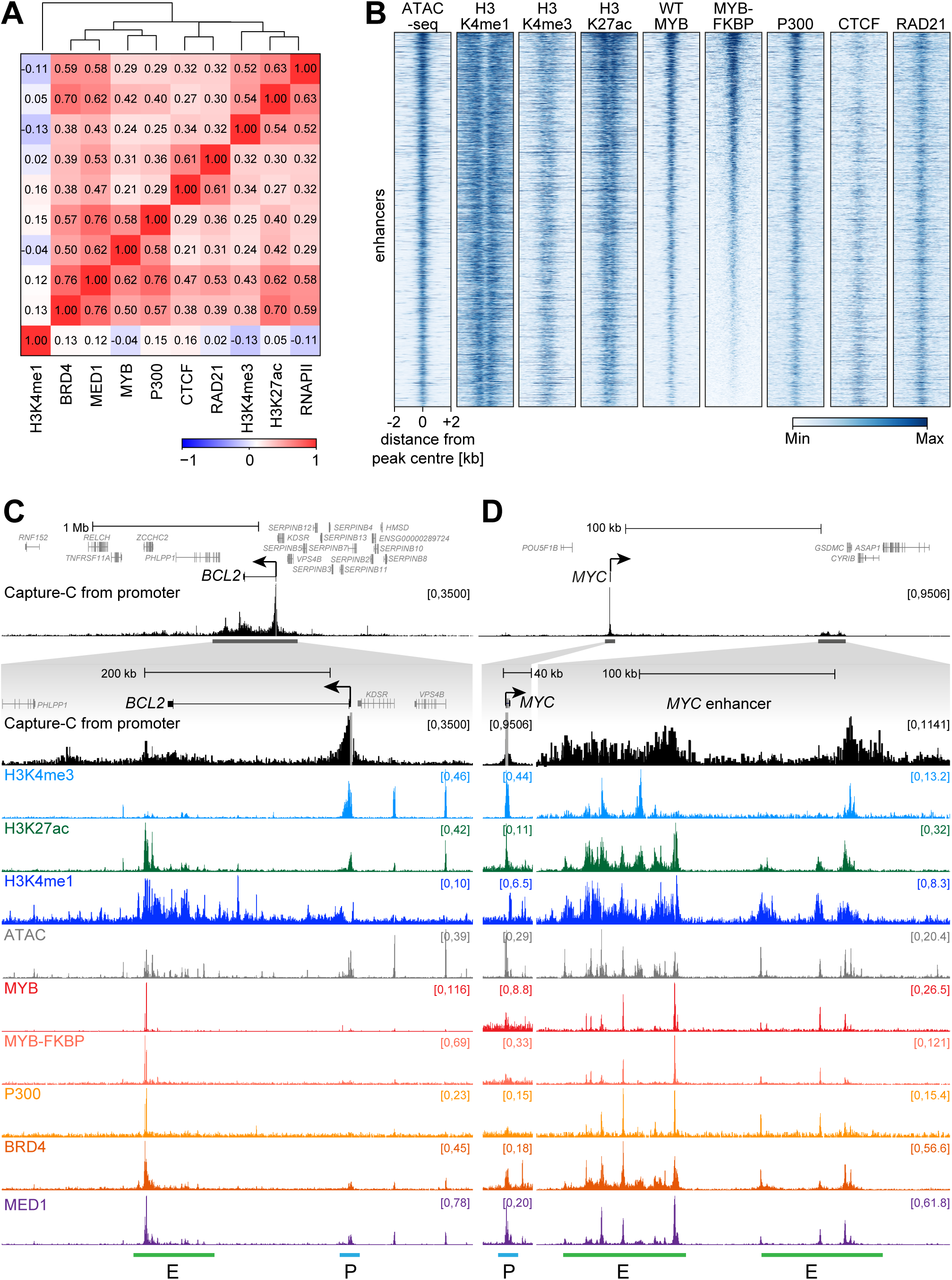
MYB binds to active enhancers in leukemia cells. (**A**) Spearman correlation coefficients for ChIP-seq data at enhancer ATAC-seq peaks in SEM cells. (**B**) Heatmap comparing levels of histone modifications and chromatin binding of MYB and co-activator proteins at enhancer ATAC-seq peaks in SEM cells. Peaks are ranked based on the relative level of wild-type MYB (WT MYB). (**C**) Capture-C, ChIP-seq and ATAC-seq at the *BCL2* gene and enhancer region (highlighted by green line labelled *E*) in SEM cells. Capture-C shows the frequency of interactions with the *BCL2* promoter (blue horizontal line labelled *P*), indicated by the vertical gray bar, and is displayed as the mean of three biological replicates. (**D**) Capture-C, ChIP-seq, and ATAC-seq data at *MYC* as in (C).

### MYB is required for the maintenance of enhancer signatures in leukemia cells

To test the requirement for MYB at enhancers, we engineered SEM cells to tag the endogenous copies of MYB with the FKBP12^F36V^ domain (*38*). Treatment with dTAG-13 induced rapid reduction in MYB protein levels (Fig. 2A) and MYB chromatin binding (Fig. S2A) from as early as 2 h post-treatment. Consistent with the key oncogenic role of MYB, dTAG-13 treatment resulted in loss of leukemia cell growth (Fig. S2B) and colony forming potential (Fig. S2C, D). Nascent *BCL2* transcript levels began to decrease after 2 h, showing high dependency on MYB, although the reduction in *MYC* expression was more delayed (Fig. S2E). Therefore, we chose a 24 h dTAG-13 treatment time to explore MYB function at enhancers.

**Fig. 2.**
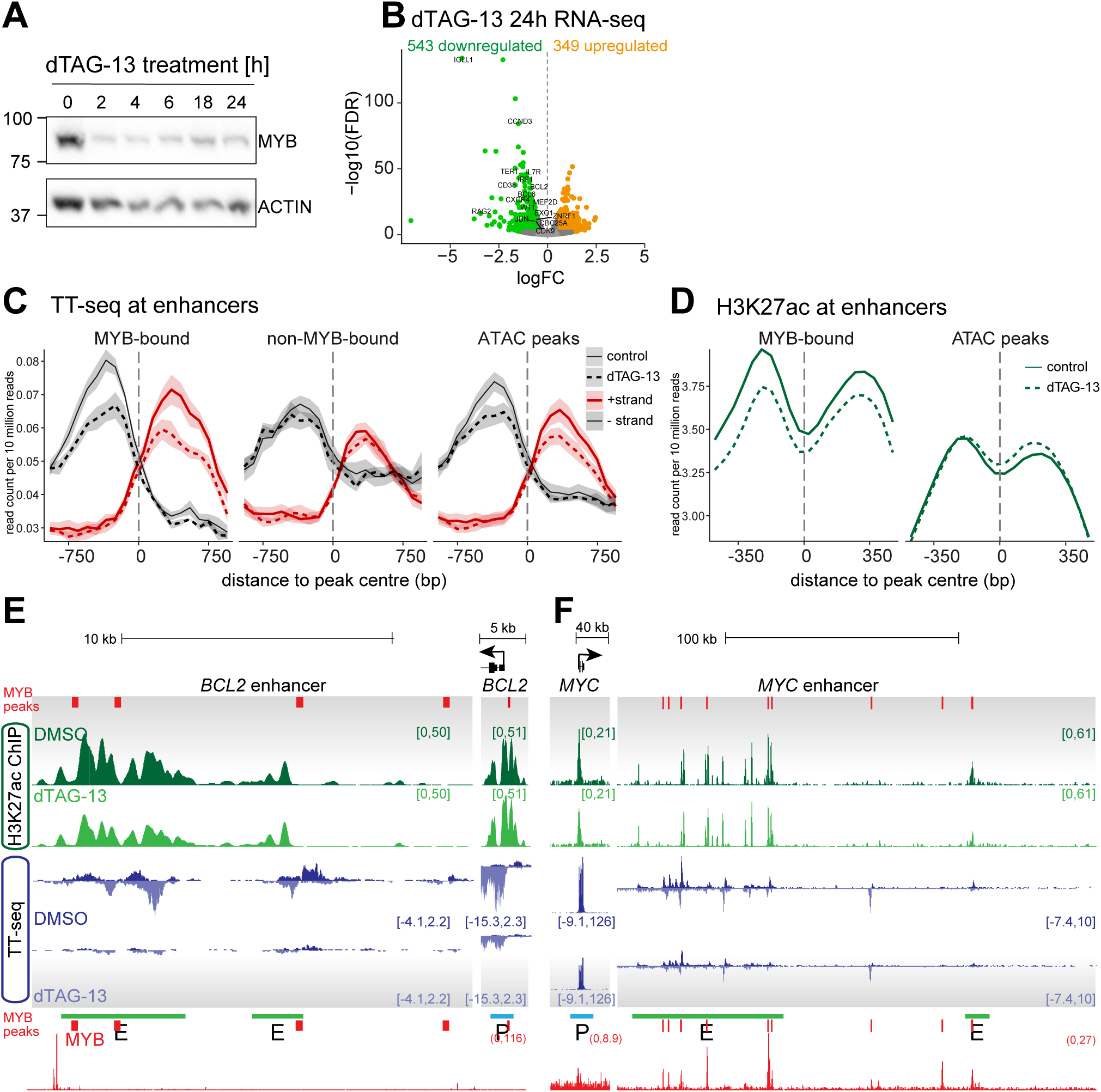
MYB is required for the maintenance of enhancer signatures in leukemia cells. (**A**) Western blot for MYB or ACTIN in MYB-FKBP12^F36V^ tagged SEM cells following addition of 0.5 μM dTAG-13. Representative of three biological replicates. (**B**) Changes in TT-seq RNA levels following 24 h treatment with dTAG-13 in MYB-FKBP12^F36V^ tagged SEM cells. Statistically significant differences (green: decreased; orange: increased; gray: unchanged) from three biological replicates, FDR < 0.05. (**C**) Strand-specific TT-seq (enhancer RNA; right) levels at intergenic MYB-bound and non-MYB-bound enhancers and all integenic enhancer ATAC-seq peaks, in control (untreated) and 24 h dTAG-13-treated MYB-FKBP12^F36V^ tagged SEM cells. Lines represent mean, shading represents ± SEM, n = 3 independent experiments. (**D**) Mean distribution of H3K27ac at MYB-bound and all enhancer ATAC peaks in control (untreated) and 24 h dTAG-13-treated MYB-FKBP12^F36V^ tagged SEM cells. H3K27ac ChIP-seq and TT-seq at (**E**) *BCL2* and (**F**) *MYC* in MYB-FKBP12^F36V^ tagged SEM cells, with or without the addition of dTAG-13 for 24 h. MYB peaks are annotated in red.

Using transient transcriptome sequencing (TT-seq) (*39*), we observed widespread changes in gene expression following 24 h MYB degradation (Fig. 2B and S2F), including downregulation of known MYB target genes *BCL2* and *MYC*, as well as *LMO4* and *CDK6* (Fig. S2G). At MYB-bound enhancers, we saw a dramatic reduction in eRNA transcription (Fig. 2C), matched with a similar decrease in H3K27ac (Fig. 2D), suggesting a direct dependency on MYB for enhancer function. *BCL2* and *MYC* both have well-characterized enhancers in SEM cells (*20, 40*), and loss of MYB enhancer binding (Fig. S2A) was associated with dramatic reductions in eRNA transcription and mnore subtle impacts on H3K27ac (Fig. 2E, F) at these MYB target genes. Taken together, this suggests that MYB may be essential for the regulation of enhancer activity.

### MYB is necessary for the maintenance of enhancer-promoter interactions at endogenous loci

Given the loss of enhancer activity associated with MYB degradation, we used the high-resolution 3C technique MCC (*23*) to ask whether MYB contributes to the spatial co-localisation of enhancers and promoters. This revealed the presence of sharp, high intensity enhancer-promoter interaction peaks (MCC peaks), which were not visible when using the lower-resolution Capture-C (Fig. 3A, B, S3A). For example, we have previously shown that the oncogene *FLT3* is regulated by a broad KMT2A-AFF1-dependent enhancer within the neighbouring *PAN3* gene (*41*). Compared to the broad interaction profile seen with Capture-C, it was striking to observe two prominent MCC peaks located at the 3’ end of this enhancer, one of which closely overlaps with a MYB binding site (Fig. 3B). We made similar observations at the enhancers of *BCL2* and *MYC*, where MCC identified MYB-bound promoter-interacting peaks (Fig. 3A, S3A). While not all promoter-interacting sites were bound by MYB, MYB binding within enhancers often co-localised with MCC peaks. Across a panel of 43 genes, we found an overlap between MYB binding and MCC peaks at most genes analyzed (Fig. S3B).

**Fig. 3.**
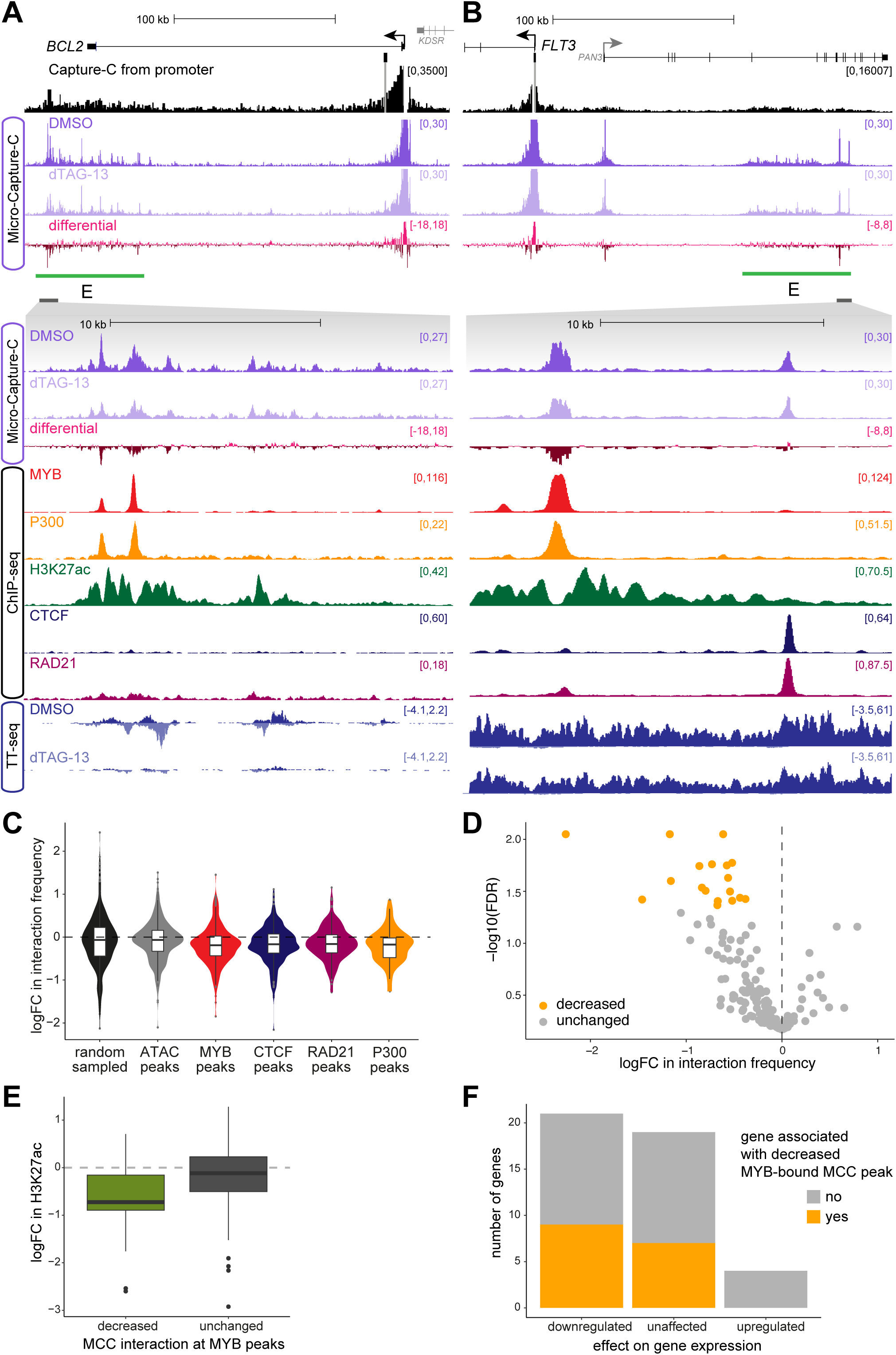
MYB is necessary for the maintenance of enhancer-promoter interactions at endogenous loci. Capture-C and ChIP-seq in SEM cells, and Micro-Capture-C (MCC) and TT-seq in MYB-FKBP12^F36V^ tagged SEM cells, with or without the addition of dTAG-13 for 24 h at the (**A**) *BCL2* and (**B**) *FLT3* loci. Enhancer regions are highlighted by the green line labelled E. Capture-C and MCC traces are scaled to emphasize distal interactions. Differential track shows the difference in interaction frequency between DMSO and dTAG-13 conditions. (**C**) Change in MCC interaction frequency (mean logFC of three replicates) between promoters and ATAC, MYB, CTCF, RAD21 and P300 peaks within approx. 500 kb of the capture point following the addition of dTAG-13 for 24 h. Violin plot shows frequency distribution. Box plot midline shows median, with upper and lower hinges showing 25th and 75th percentile, respectively. Upper and lower hinges extend to the largest and smallest datapoints within 1.5 times the interquartile range of either hinge. (**D**) Changes in MCC interaction frequency at MYB peaks following 24 h treatment with dTAG-13 in MYB-FKBP12^F36V^ tagged SEM cells. Statistically significant differences (yellow: decreased; gray: unchanged) from three biological replicates, FDR < 0.05. (**E**) H3K27ac levels at MYB peaks associated with with decreased or unchanged MCC interaction. (**F**) Correlation of loss of MCC interaction at MYB peak with changes in gene expression.

MYB degradation with dTAG-13 had little or no effect on the frequency of enhancer-promoter association for the majority of MCC peaks (Fig. S3C). However, by focussing on MYB-bound MCC peaks, we observed a strong bias towards decreased interaction frequency (Fig. 3C). Statistically significant peaks were exclusively associated with a loss of interaction (Fig. 3D). Significantly decreased MCC peaks were also enriched for key enhancer proteins P300, BRD4 and MED1 (Fig. S3D), revealing a role for MYB in the maintenance of enhancer-promoter interactions. In support of this, there was a clear correlation between decreases in interaction frequency and decreased H3K27ac following dTAG-13 treatment (Fig. 3E, S2E, F). Finally, of the genes that showed a loss of promoter interactions with MYB-bound MCC peaks, the majority were downregulated following MYB degradation, indicating the transcriptional consequences of loss of MYB binding (Fig. 3F).

These features are clearly demonstrated at the MYB target gene *BCL2*. At the *BCL2* enhancer, the sites that show major decreases in promoter contact are bound by MYB (Fig. 3A, see MCC differential track), and also show reduced H3K27ac and eRNA transcription (Fig. 2E, 3A). Thus, the presence of MYB is required for localised contact with the *BCL2* promoter, along with enhancer activity and gene expression. Similarly, at the *FLT3* and *MYC* enhancers, there was a specific loss of promoter interaction, associated with decreased H3K27ac and eRNA transcription (Figs. 3B, 2F, S3A).

Overall, these results demonstrate that MYB is required for key enhancer activities, including histone acetylation, eRNA transcription and physical proximity with the target gene promoter, the loss of which results in transcriptional downregulation. We therefore sought to ask whether these activities mean that binding of MYB alone is sufficient to generate an enhancer.

### MYB^TA^ is sufficient to initiate de novo enhancer activity

Most attempts to explore TF function have been in the context of endogenous enhancers, with additional TF binding sites present, or in exogenous constructs lacking a chromatin environment. We wanted to determine if the binding of MYB alone could generate enhancers de novo in a chromatin context. To explore this, we used a well-characterised chromatin anchoring system, based on an array of *Tet operator* (*TetO*) sequences flanked by inert human chromatin, which is inserted into mESCs (*41–43*). By fusing Myb to the Tet repressor (TetR) DNA binding domain (Fig. 4A), we could directly tether the TF to the *TetO* locus and assess its effect on the chromatin environment (Fig. 4B). Since the activity of full length MYB may be attenuated through intramolecular interactions between its N- and C-termini, we isolated the transactivation domain of mouse Myb (Myb^TA^), which is sufficient for Myb transactivating activity (*44–46*).

**Fig. 4.**
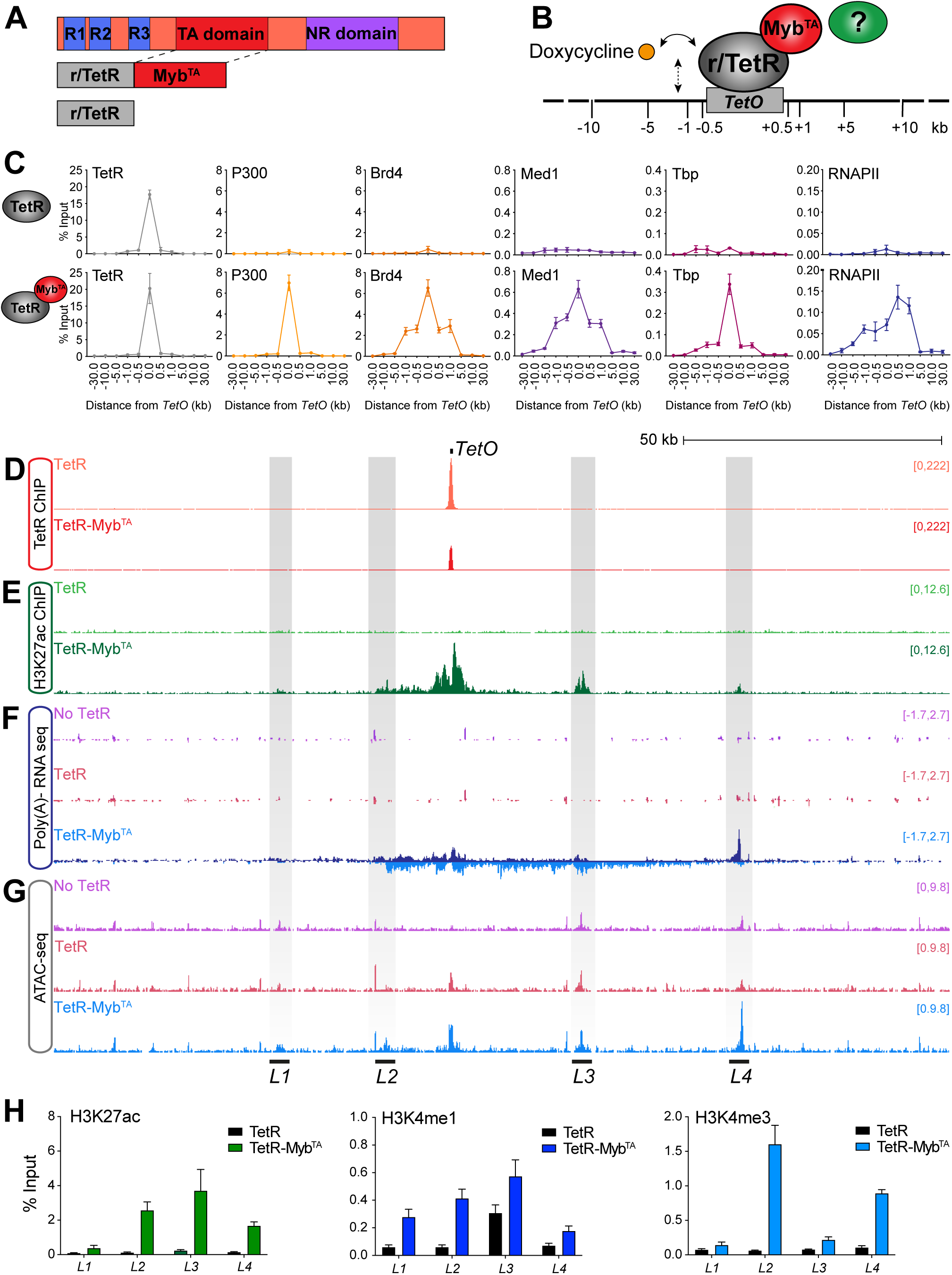
TetR-MYB^TA^ is sufficient to initiate de novo enhancer activity. (**A**) Schematic of MYB protein structure and conserved domains, with the DNA binding (DB) domain with tandem repeats R1-3, the transactivation (TA) domain and the negative regulatory (NR) domain, as well as r/TetR-Myb^TA^ and control (r/TetR only) proteins. (**B**) Targeting of factors to *TetO* via the r/TetR DNA-binding domain. Numbers represent qPCR primer positions (kb) with respect to the *TetO* array. Doxycycline disrupts DNA binding of TetR but induces binding of rTetR. (**C**) ChIP-qPCR analysis across the *TetO*-containing locus in cell lines stably expressing either TetR (top row) or TetR-Myb^TA^ (bottom row), using antibodies against TetR, p300, Brd4, Tbp, Med1 and RNAPII. Error bars represent the standard deviation from three biological replicates. (**D**) TetR ChIP-seq, (**E**) H3K27ac ChIP-seq, (**F**) Poly-A minus RNA-seq and (**G**) ATAC-seq at the integrated *TetO* array on mouse chromosome 8. *L1-4* are highlighted in gray. (**H**) ChIP-qPCR for H3K27ac (left), H3K4me1 (middle) and H3K4me3 (right) at *L1-4* in TetR (black bars) or TetR-Myb^TA^ (colored bars) expressing cells. Error bars represent the standard deviation from three biological replicates.

Binding of both TetR-Myb^TA^ and the control TetR was restricted to the *TetO* locus (Fig. 4C and D). TetR-Myb^TA^ recruited its known co-activator, p300 (Fig. 4C), resulting in enrichment of H3K27ac at the *TetO* locus, in a region previously devoid of this modification (Fig. 4E, S4A). Additionally, we found a higher ratio of H3K4me1 to H3K4me3 enrichment at *TetO*, which is a hallmark of enhancer elements (*6, 47*) (Fig. S4A), and localisation of key enhancer-associated proteins Brd4 and Mediator (Fig. 4C, S4B). We also saw TetR-Myb^TA^ dependent binding of Tbp and RNAPII, suggesting the potential for transcription initiation (Fig. 4C, S4B). Together, this indicates that binding of TetR-Myb^TA^ at the *TetO* locus had generated an enhancer-like element.

To determine if this enhancer-like element was being actively transcribed, we performed RNA-seq of polyA-depleted transcripts to identify nascent transcription. The ∼150 kb region centered on *TetO* was transcriptionally silent in the absence of TetR-Myb^TA^ binding. Remarkably, in cells expressing TetR-Myb^TA^, we detected transcription on both strands throughout this locus. While some of this may have initiated at *TetO* itself, much appeared to originate at other sites, up to 50 kb away from the TetR-Myb^TA^ binding site (Fig. 4F).

We next used ATAC-seq to identify accessible regions that could be sites of transcription initiation. While some ATAC-seq peaks were visible in cells expressing no fusion protein or TetR alone (annotated *L1*, *L3* and *L4*), accessibility at these peaks increased upon expression of TetR-Myb^TA^, along with the appearance of a novel ATAC-seq peak (annotated *L2*) (Fig. 4G). Strikingly, unique to TetR-Myb^TA^ expressing cells, we observed H3K27ac enrichment at all four loci, suggesting that these distal regions were activated by TetR-Myb^TA^ binding at *TetO* (Fig. 4E, H, S4C). Of note, *L1* and *L3* appeared to be origins of bidirectional transcription, consistent with enhancer-like activity, while transcription from *L2* and *L4* was directionally biased, reminiscent of transcription start sites at promoters (Fig. 4F, S4C). Consistent with this, *L1* and *L3* were enriched for the enhancer mark H3K4me1, whereas *L2* and *L4* were associated with the promoter modification H3K4me3 (Fig. 4H). These modifications were specifically induced in TetR-Myb^TA^ expressing cells. Taken together, these data strongly argue that the binding of TetR-Myb^TA^ to chromatin in isolation establishes an enhancer-like regulatory element capable of activating distant cryptic promoters and enhancers.

### Myb^TA^ binding induces – and is necessary for – chromatin looping

To understand the kinetics of TetR-Myb^TA^-dependent enhancer activation, we exploited the properties of the Tet-On and Tet-Off tetracycline-controlled gene expression systems (*48*). On addition of doxycycline, binding of TetR-Myb^TA^ to *TetO* is disrupted, whereas rTetR-Myb^TA^ only binds in the presence of doxycycline. We therefore used rTetR-Myb^TA^ expressing mESCs to explore the induction of enhancer activity. Addition of doxycycline resulted in the rapid induction of transcription from the *TetO* locus as well as the distal loci *L1-4* within 30 min, with further increases at 24 h (Fig. 5A). Of interest, we noted that the onset of transcription appeared to temporally precede H3K27ac deposition (Fig. 5A, S5A), suggesting that appreciable levels of H3K27ac may be unnecessary for the induction of transcription initiation.

**Fig. 5.**
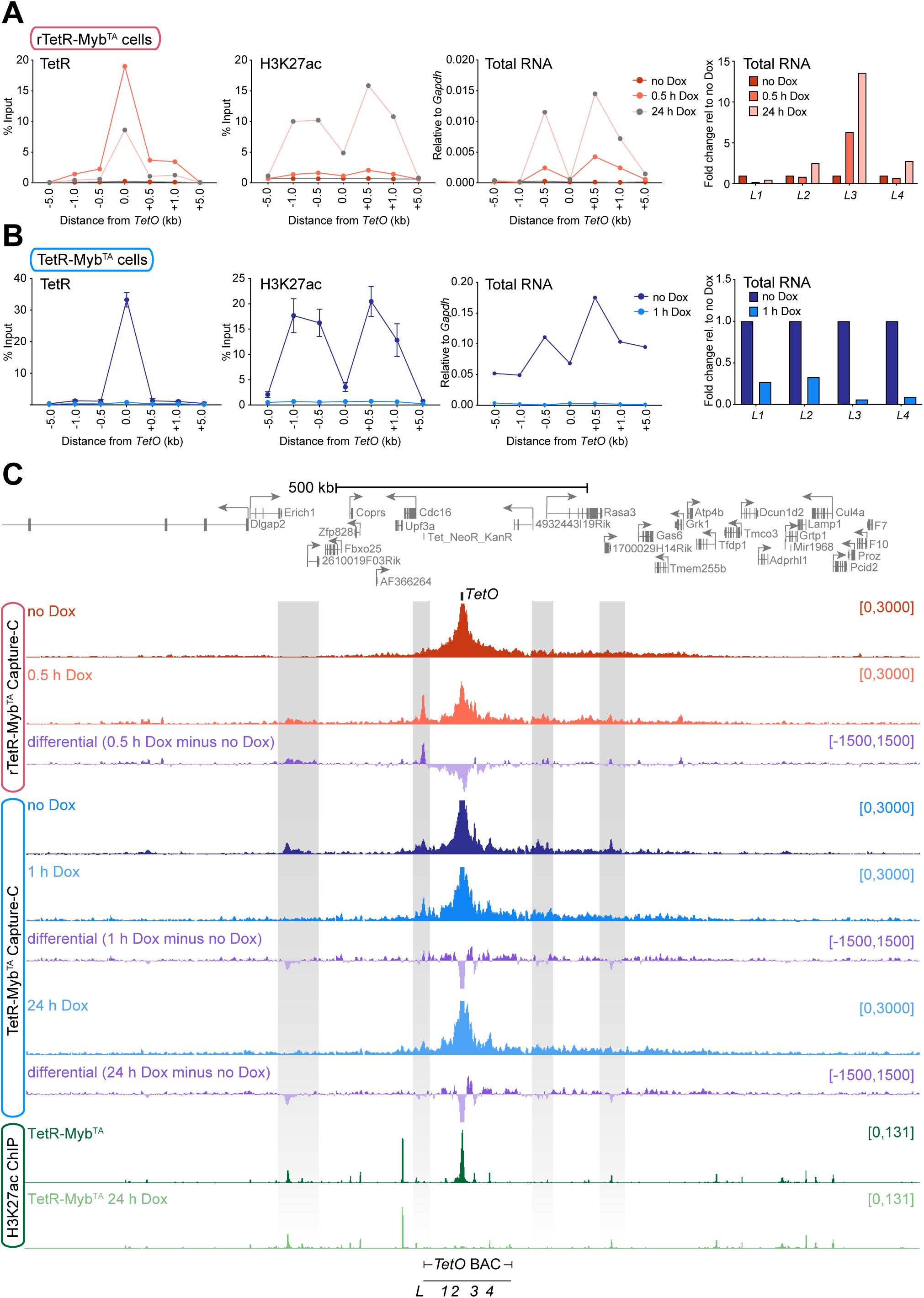
MYB^TA^ binding induces – and is necessary for – chromatin looping. (**A**) ChIP-qPCR analysis in rTetR-Myb^TA^ expressing cells for r/TetR (1^st^ panel) and H3K27ac (2^nd^ panel) and qRT-PCR (3^rd^ panel) across the *TetO* containing locus and at *L1-4* (4^th^ panel) following doxycycline treatment for 0.5 and 24 h. Error bars represent the standard deviation from three biological replicates. (**B**) As in (A), ChIP-qPCR and qRT-PCR in TetR-Myb^TA^ expressing cells following doxycycline treatment for 1 h. (**C**) Capture-C and H3K27ac ChIP-seq in untreated and doxycycline-treated rTetR-Myb^TA^ and TetR-Myb^TA^ expressing cells across the *TetO* locus. Differential track shows the difference in interaction frequency between untreated and doxycycline conditions.

We used doxycycline treatment in TetR-Myb^TA^ cells to test the requirement for TetR-Myb^TA^ in enhancer maintenance. A short (1 h) treatment with doxycycline, which abrogated TetR-Myb^TA^ binding, resulted in a dramatic loss of both transcription and H3K27ac from *TetO* and the distal loci (Fig. 5B, S5B). Thus, both local and distal enhancer activity is absolutely dependent on the continued presence of TetR-Myb^TA^ at *TetO*.

We probed the role of p300/Cbp in directing transcription at this locus using Myb^TA^ mutants that partially (M303V) or fully (L302A) disrupt binding to p300/Cbp (*30, 49, 50*). TetR-Myb^TA^ L302A localised to *TetO*, but did not result in binding of p300 at the locus and was unable to induce detectable enrichment of H3K27ac or transcription at *TetO* or the distal loci (Fig. S5C, D). In contrast, use of TetR-Myb^TA^ M303V partially reduced, but did not eliminate, binding of p300 and H3K27ac deposition, compared to wild-type TetR-Myb^TA^ (Fig. S5C). Strikingly, however, transcription was much more dramatically reduced, to approx. 1% of that induced by TetR-Myb^TA^ (Fig. S5D). This indicates that p300/Cbp alone may not be sufficient to induce transcription at this enhancer-like element, and other factors may also be required.

For TetR-Myb^TA^ to activate acetylation and transcription at distal loci, these regions may need to come into close physical proximity with *TetO*. Given the effect of MYB degradation on enhancer-promoter interactions in SEM cells, we hypothesised that Myb binding may induce contact between *TetO* and distal chromatin. To test this, we performed Capture-C (*19*, *20*) in rTetR-Myb^TA^ cells, using a *TetO* specific capture probe, to measure interactions with the TetR-Myb^TA^ binding site (Fig. 5C).

In the absence of rTetR-MYB^TA^ binding (no Dox) we observed a normal decay in interaction frequency on either side of *TetO* (Fig. 5C) consistent with random chromatin interactions. However, within 30 min of doxycycline-induced binding of TetR-Myb^TA^ at *TetO*, we observed the appearance of chromatin interactions at the distal loci *L1, L3* and *L4* (Fig. S5E). *L2* is only 14.5 kb from the *TetO*, so our ability to detect increased interaction with the *TetO* may be obscured by proximity to the capture locus. Most strikingly, in addition to the appearance of these interactions within the inserted *TetO* BAC sequence, we also detected longer range interactions up to 400 kb away from *TetO* (highlighted in gray, Fig. 5C). These interactions were also detected in the TetR-Myb^TA^ expressing cell line. In confirmation of our endogenous

MYB degradation experiments (Fig. 3), many of these interactions were lost following TetR-Myb^TA^ displacement upon doxycycline treatment, arguing that Myb is responsible for establishing, and required to maintain, chromatin interactions (Fig. 5C, S5E). This is particularly striking at the *L3* locus (Fig. S5E), which is marked with high levels of TetR-MYB^TA^-dependent H3K27ac and H3K4me1 (Fig. 4E, H). Taken together, our data argue that MYB is capable of remodelling large chromatin domains, driving productive interactions to induce transcription at distal loci.

## Discussion

Here, we show that MYB binding is essential for transcription and drives discrete enhancer-promoter contacts. Results using high-resolution 3C methods have previously suggested that enhancer-promoter contact points are precisely localized to TF binding sites (*23–25*), and recent work with LDB1 suggests that TF/cofactor interactions themselves can be essential for widespread loop formation (*17*). It is unclear if the ability to form loops is a general feature of TF activity, or if it occurs at only a subset of locations. In our work, we find that MYB degradation results in the loss of many interaction sites that are precisely localised over MYB-bound loci, suggesting that it is MYB itself (or the co-activators it recruits) that drives enhancer-promoter contact in these specific regions.

From our work, we note two points. Firstly, we only observed this effect at a small minority of MYB binding sites; at many MYB-bound enhancers, degradation of MYB had little or no effect on the interaction with the promoter, indicating that other factors contribute at these loci. This suggests that an essential requirement for MYB is highly context specific, perhaps reflecting the absence of alternative, redundant factors. Secondly, in those cases where MYB degradation reduced the punctate interaction at the MYB peak, the broader interaction profile of the enhancer remained intact, and overall the enhancer continued to interact with the promoter in the absence of MYB. Thus at most enhancers, it is not a single TF driving the interaction profile, but instead the sum of interactions across multiple protein complexes. Even where MYB is required, adjacent interactions within the enhancer are maintained by alternative mechanisms. Future work will determine if this activity is the main function of a handful of key TFs or cofactors such as MYB and LDB1, or of all TFs contribute to enhancer-promoter crosstalk in this way.

To understand the role of MYB in the absence of adjacent TF binding events, we anchored Myb in a gene desert region. We found that the Myb activation domain alone has the ability to recruit co-activators and induce long-range interactions, activating cryptic promoters, and that the continued presence of Myb is required to maintain many of these interactions. This suggests that mediating long-range gene activation is an intrinsic property of the MYB protein. How Myb selects sites for activation is unclear. However, we observed low-level chromatin accessibility in the absence of TetR-Myb^TA^, indicating the potential for these sites to be bound by other TFs, poised for activation upon interaction with Myb. It seems likely that this de novo activity is context dependent, as regulatory elements can retain function when removed from their endogenous context, but are still responsive to the overall cell environment (*51*).

MYB-dependent interactions between the enhancer and promoter are likely driven by MYB co-activator recruitment, and subsequent stabilisation of the RNAPII complex. MYB is known to function via interaction with P300/CBP (*30–32*), so this may provide a mechanism for enhancer-promoter interactions. Indeed, histone acetylation and 3D contacts were both rapidly lost following TetR-Myb^TA^ dissociation from chromatin. Further, upon MYB degradation, we found a correlation between reductions in H3K27ac and interaction frequency at MYB-bound enhancer-promoter contact peaks, although it is unclear whether this reflects two independent activities of MYB, or if the two are interdependent. By using TetR-Myb^TA^ constructs with mutations that disrupt its interaction with p300/Cbp (*30, 49, 50*), we observed disruption of both histone acetylation and p300/Cbp recruitment at *TetO* and transcription at distal loci. Strikingly, the effect on transcription was much more dramatic, indicating that there may be additional, acetyltransferase-independent functions of p300/Cbp, or additional cofactors that interact with the Myb TA domain, contributing to activation of distal loci. Whether the mechanisms that facilitate enhancer-promoter contact and promoter activation are the same or distinct remains to be established. We have previously found that degradation of the enhancer-associated factor BRD4 disrupts transcription without affecting 3D genome interactions (*20*), so there is no intrinsic requirement for the two activities to be driven by the same process. In our system, MYB appears sufficient to both drive interactions with distal loci and promote transcriptional activation at these sites. Further work is required to explore how these activities are mediated.

## Materials and Methods

### Cell culture

SEM cells were purchased from DSMZ and cultured in Iscove’s Modified Dulbecco’s Medium (IMDM) supplemented with 10% fetal bovine serum (FBS, ThermoFisher Scientific) and 2 mM GlutaMAX (ThermoFisher Scientific). Mouse ESCs with a *TetO* array insertion (TOT2N mESC) were the kind gift of Prof. Rob Klose (University of Oxford) and were grown in Dulbecco’s Modified Eagle’s Medium (DMEM) supplemented with 10% FBS, non-essential amino acids (ThermoFisher Scientific), GlutaMAX, LIF and β-mercaptoethanol. All cell lines were confirmed free from mycoplasma contamination. Colony forming assays were conducted by plating 500 cells mixed with 1 ml H4100 Methylcellulose medium (StemCell Technologies) in a 3 cm diameter dish in triplicate, then allowing to grow for 14 days before counting.

### TetR cell lines

The Myb transcriptional activation domain (Myb^TA^) and Myb^TA^ mutant sequences (Myb^TA^ L302A and Myb^TA^ M303V) were cloned into the original pCAGFS2TetR vector, downstream of the FS2-TetR open reading frame. The rTetR-MYB^TA^ construct was generated by dsODN exchange cloning, replacing the TetR domain with the rTetR sequence (*52*). Plasmids were transfected into TOT2N mESCs with Lipofectamine-2000, and clones stably expressing the transfected construct were selected using 1 µg/ml puromycin, then confirmed by western blotting. As a negative control, parental mESCs (no TetR) or cells expressing the TetR domain alone were used. Where indicated, cells were treated with 1 µg/ml doxycycline.

### Generation of degron cell lines

SEM cells were edited to incorporate *FKBP12^F36V^-HA-P2A-mNeonGreen*, immediately prior to the stop codon of both alleles of *MYB*, by CRISPR/Cas9-mediated homology-directed repair (HDR), as previously described (*13*). Cells were co-electroporated with pX458 (encoding *Cas9*, one of three sgRNA sequences targeting the end of *MYB* (Supplementary Table S1), and *mRuby*) and an HDR construct containing the *FKBP12^F36V^-HA-P2A-mNeonGreen* sequence flanked by 500 bp regions with homology to either side of the *MYB* stop codon. Fluorescence-activated cell sorting was used to isolate mRuby-positive cells after 24h, and cells were allowed to grow for 1-2 weeks, after which cells were sorted again. mNeonGreen-positive cells were plated onto H4100 Methylcellulose medium and after 1-2 weeks, individual colonies screened for homozygous editing of the *MYB* gene by PCR and western blotting.

### Chromatin immunoprecipitation (ChIP)

ChIP was conducted as described previously (*43, 53*). Briefly, 10^7^ cells were fixed for histone modifications (1% formaldehyde for 10 min) or TFs/cofactors (2 mM disuccinimidyl glutarate for 30 min, then 1% formaldehyde for 30 min). Fixed cell pellets were lysed in 120 µl lysis buffer (10 mM Tris-HCl pH 8.0, 1 mM EDTA, 1% SDS) and sonicated with an ME220 sonicator (Covaris) to generate 200-300 bp fragments. After pelleting insoluble material, the supernatant was diluted 10x, then 5 µl protein A- and G-coupled dynabeads (ThermoFisher Scientific) were added to pre-clear chromatin prior to immunoprecipitation. An input sample was taken, then 2 µg antibody (Supplementary Table S2) was added and the sample was incubated overnight with rotation at 4 °C. Antibody-chromatin complexes were isolated by addition of 15 µl protein A- and G-coupled dynabeads, after which the beads were washed 3x with RIPA buffer (50mM HEPES-KOH (pH 7.6), 500mM LiCl, 1mM EDTA, 1% NP40 and 0.7% sodium deoxycholate) and once with Tris-EDTA. Samples were eluted with lysis buffer, then RNase A- and proteinase K-treated and crosslinks were reversed by overnight incubation at 65 °C. DNA was purified using the QIAquick PCR Purification Kit (Qiagen) and quantified by qPCR, normalized to input. Primer sequences are detailed in Supplementary Table S1. ChIP-seq libraries were generated with the NEBNext Ultra II DNA library prep kit (NEB), then sequenced by 75 cycle paired-end sequencing on a NextSeq machine (Illumina).

### ATAC-seq

ATAC-seq was conducted using 50000 live cells using Nextera Tn5 transposase (Illumina) as previously described (*54*), and sequenced by 75 cycle paired-end sequencing on a NextSeq machine (Illumina).

### ChIP-seq and ATAC-seq analysis

FASTQ file quality was confirmed by FastQC (v0.12.1; http://www.bioinformatics.babraham.ac.uk/projects/fastqc/), after which reads were trimmed using trim_galore with Cutadapt (v0.6.10; https://www.bioinformatics.babraham.ac.uk/projects/trim_galore/). Trimmed reads were mapped to the hg38 reference genome using Bowtie 2 (v2.5.1) (*55*). PCR duplicates were removed with picard MarkDuplicates (v3.0.0; http://broadinstitute.github.io/picard). The ENCODE Blacklist (https://doi.org/10.1038/s41598-019-45839-z) was used to remove problematic mapping regions from the aligned files and further QC of the aligned files was performed using samtools (v1.17) (*56*). Peaks were called and bigwigs were generated using HOMER (v4.11) (*57*) and visualized in the UCSC genome browser (*58*). Putative enhancers were identified by the intersection of non-promoter ATAC-seq peaks with H3K27ac peaks. Metaplots were generated using the HOMER function annotatePeaks.pl (*57*), centered on enhancer ATAC-seq peaks. Heatmaps were generated using deepTools (v3.5.1) (*59*).

### Transient Transcriptome sequencing (TT-seq)

TT-seq was conducted as previously described (*39*). Briefly, spike-in RNA was generated by in vitro transcription of exogenous plasmid sequences in the presence of 4S-UTP (Jena Bioscience), using the MEGAscript kit (ThermoFisher Scientific). 5×10^7^ SEM cells were treated with 500 µM 4-thiouridine for 5 min, before RNA isolation by Trizol extraction (ThermoFisher Scientific) with the addition of 60 ng spike-in RNA, then DNase I-treatment. Labelled nascent RNA was fragmented briefly by sonication (Covaris), biotinylated with EZ-link biotin-HPDP (ThermoFisher Scientific), then purified by streptavidin bead pull-down (Miltenyi). Strand-specific libraries were prepared using the NEBNext Ultra II Directional RNA Library Prep Kit for Illumina (NEB) and sequenced by 150 cycle paired-end sequencing on a NextSeq machine (Illumina).

### PolyA-minus RNA sequencing

RNA was extracted from cells using the RNeasy Mini Kit (Qiagen). PolyA-minus RNA was isolated using the NEBNext Poly(A) mRNA magnetic isolation module (retaining the unbound fraction), then used to generate a strand-specific library using the NEBNext Ultra II Directional RNA Library Prep Kit for Illumina (NEB). Libraries were sequenced by 150 cycle paired-end sequencing on a NextSeq machine (Illumina).

### RNA-seq analysis

Reads were subjected to quality checking by fastQC (v0.12.1) and trimming using trim_galore (v0.6.10) to remove contaminating sequencing adapters, poor quality reads and reads shorter than 21 bp. Reads were then aligned to hg38 using STAR (v2.4.2a) (*60*) in paired-end mode using default parameters. Gene expression levels were quantified as read counts using the featureCounts function from the Subread package (v2.0.2) (*61*) with default parameters. The read counts were used to identify differential gene expression between conditions using the EdgeR (v3.12) (*62*) package. For TT-seq, spike-in RNA levels were quantified by mapping to a custom genome using featureCounts and used to normalize the output of EdgeR.

### qRT-PCR

RNA was extracted from cells using the RNeasy Mini Kit (Qiagen), then reverse transcribed using SuperScript III (ThermoFisher Scientific) with random hexamer primers. cDNA was analyzed by qPCR using Taqman probes (ThermoFisher Scientific) or SyBr Green (ThermoFisher Scientific). The housekeeping genes *GAPDH* and *YWHAZ* were used for normalization. Taqman probe IDs and SyBr primer sequences are detailed in Supplementary Table S1.

### Western blotting

Salt-soluble proteins were extracted from PBS-washed cells using high-salt lysis buffer (20 mM Tris-HCl pH 8.0, 300 mM KCl, 5 mM EDTA, 20 % glycerol, 0.5 % IGEPAL CA-630, protease inhibitor cocktail). Proteins were separated by SDS-PAGE, transferred to PVDF membrane, then probed (Supplementary Table S2) and visualized using enhanced chemiluminescence.

### Next Generation Capture-C

Capture-C was conducted as described previously (*37, 63*), using 2×10^7^ cells per replicate. Briefly, DpnII-generated 3C libraries were sonicated to 200 bp fragments and Illumina paired-end sequencing adaptors (New England Biolabs) were added using Herculase II (Agilent). Indexing was performed in duplicate to maintain library complexity, with libraries pooled after indexing. Enrichment was performed using biotinylated Capture-C probes (Supplementary Table S3) (*53*), with two successive rounds of hybridization, streptavidin bead pulldown (ThermoFisher Scientific), bead washes and PCR amplification using the HyperCapture Target Enrichment Kit (Roche). Samples were sequenced by paired-end sequencing with a 300 cycle high-output Nextseq 500 kit (Illumina). Data analysis was performed using CapCruncher v0.2.0 (*63*) (https://doi.org/10.5281/zenodo.6326102) and statistical analysis was performed as described (*37, 63*).

### Micro-Capture-C

Micro-Capture-C was performed as described (*23, 64*). Briefly, 10^7^ SEM cells were fixed with 2% formaldehyde for 10 min, then quenched with glycine and PBS-washed. After permeabilization with 0.005% digitonin for 15 min, cells were snap frozen. Thawed cells were pelleted and resuspended in reduced-calcium MNase buffer (10 mM Tris-HCl pH7.5, 1 mM CaCl_2_), then divided into three aliquots. Cells were titrated with different micrococcal nuclease (NEB) concentrations for 1h at 37 °C with shaking at 550 rpm. The reaction was stopped by addition of EGTA to 5 mM, with 200 µl removed to assess digestion. Remaining cells were pelleted and washed with PBS/EGTA, then cell pellets were resuspended in DNA ligase buffer, supplemented with 400 µM dNTPs and 5 mM EDTA, before addition of DNA Polymerase I large (Klenow) fragment (NEB) to 100 U/µl, T4 polynucleotide kinase (NEB) to 200 U/µl and T4 DNA ligase (Thermo Scientific) to 300 U/µl. The reaction was incubated for 2h at 37 °C, then for 8h at 20 °C, at 550 rpm, then cooled to 4 °C. The digested and ligated chromatin was decrosslinked at 65 °C in the presence of proteinase K, then DNA was purified using a DNeasy Blood and Tissue Kit (Qiagen). Digestion and ligation was assessed by D1000 TapeStation (Agilent). Library preparation, indexing and capture were performed as described for NG Capture-C. Probes used for capture are given in Supplementary Table S3. MCC analysis was performed using the MCC pipeline (*23*) (https://github.com/jojdavies/Micro-Capture-C). MCC peaks were called from merged bigwig tracks from each oligo viewpoint using Lanceotron (*65*), and were then filtered on peak size, width and distance from the viewpoint. Unique junctions within each peak in the control and treated conditions were counted before normalization, using the total cis-unique ligation junctions for that peak’s corresponding viewpoint (*23*). Junction counts were compared between conditions using a Student’s T-test. MCC peaks included in the analysis were intersected with SEM cell ChIP-seq peaks from in-house datasets using Pybedtools with default paramters. The enrichment score was calculated as the number of statistically skewed MCC peaks with factor bound divided by the total number of MCC peaks used for the analysis.

## Funding

Medical Research Council (MRC, UK) Molecular Haematology Unit grant MC_UU_00016/6 and MC_UU_00029/6. (T.A.M., A.L.S.)

MRC Clinical Research Training Fellowship MR/M003221/1 (I.-J.L.)

Anya Sturdy Charitable Trust for Medical Research Fellowship A130320/129280 (I.-J.L.) Kay Kendall Leukemia Fund Intermediate Fellowship KKL1443 (N.T.C.)

The Lister Institute (J.O.J.D.)

MRC Molecular Haematology Unit grant MC_UU_00029/04 (J.O.J.D.) Wellcome Trust 225220/Z/22/Z (J.O.J.D.)

Oxford National Institute of Health Research Biomedical Research Centre NIHR203311 (J.O.J.D.)

MRC Molecular Haematology Unit grant MC_UU_00029/8 (P.V.) Blood Cancer UK Programme Continuity Grant 13008 (P.V.) Cancer Research UK SEBCATP-2022/100011 (N.D.)

## Author Contributions

Conceptualization: I.-J.L., N.T.C. and T.A.M.

Methodology: I.-J.L., N.T.C. and T.A.M.

Investigation: I.-J.L., N.D., N.E.J. and N.T.C.

Formal analysis: I.-J.L., J.R.Harman., A.L.S., J.C.H. and N.T.C.

Funding acquisition: T.A.M.

Supervision: P.V., J.O.J.D., J.R.Hughes and T.A.M.

Writing – Original Draft: I.-J.L., N.T.C. and T.A.M.

Writing – Review & Editing: All authors

## Data and materials availability

All high throughput sequencing data have been deposited in the Gene Expression Omnibus (GEO) under accession number GSEXXXXXX. Cell lines are available from the corresponding authors, upon completion of a Materials Transfer Agreement.

## Supplemental Figure Legends

**Fig. S1.**
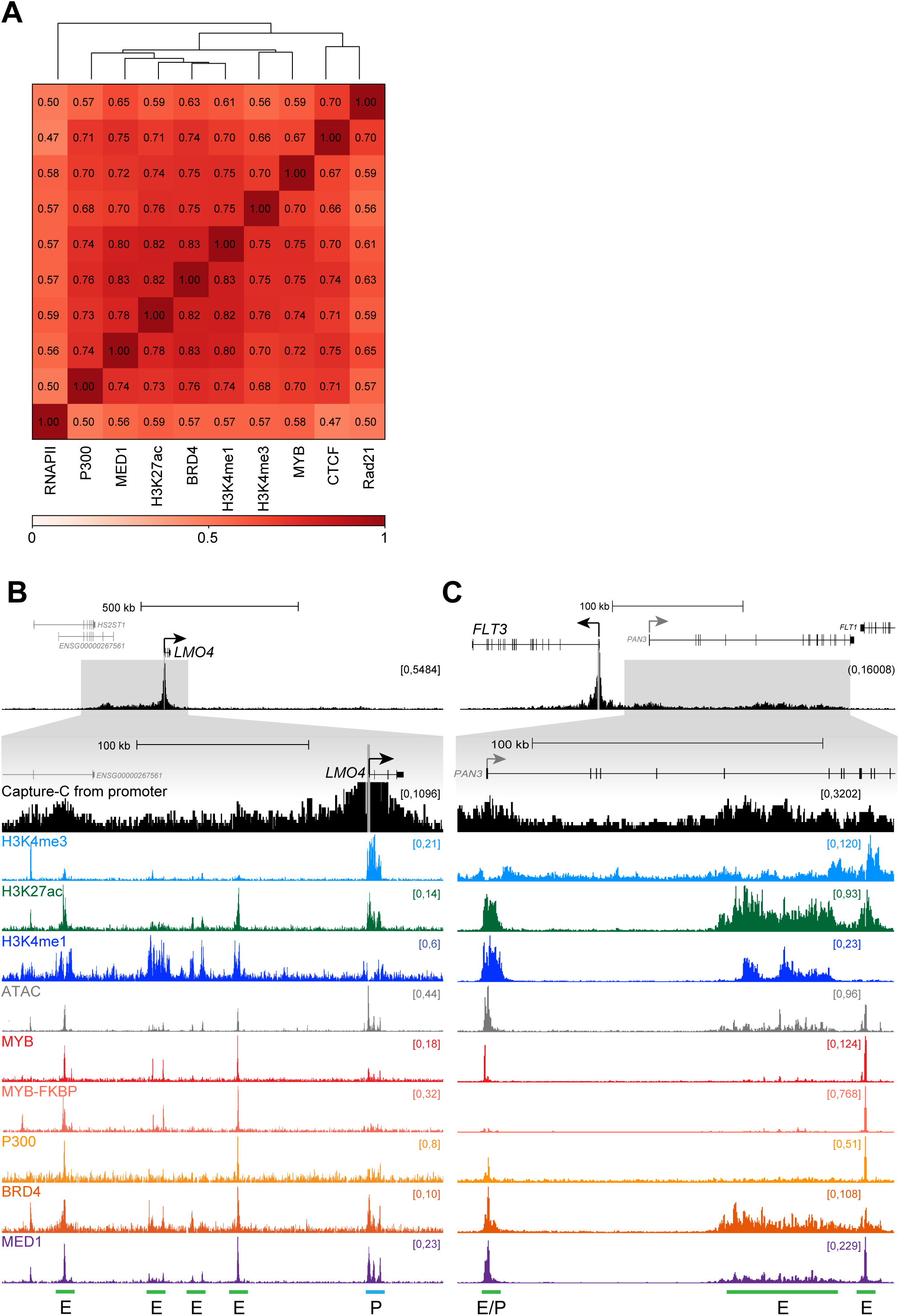
(**A**) Genome-wide Spearman correlation coefficients for ChIP-seq data at ATAC-seq peaks in SEM cells. (**B**) Capture-C, ChIP-seq and ATAC-seq at the *LMO4* gene and enhancer regions (green horizontal lines labelled *E*) in SEM cells. Capture-C was conducted using the promoter (blue horizontal line labelled *P*) as the viewpoint, indicated by the vertical gray bar, mean of three biological replicates. Capture-C traces are scaled to emphasize distal interactions (**C**) Capture-C, ChIP-seq, and ATAC-seq data at *FLT3* as in (B).

**Fig. S2.**
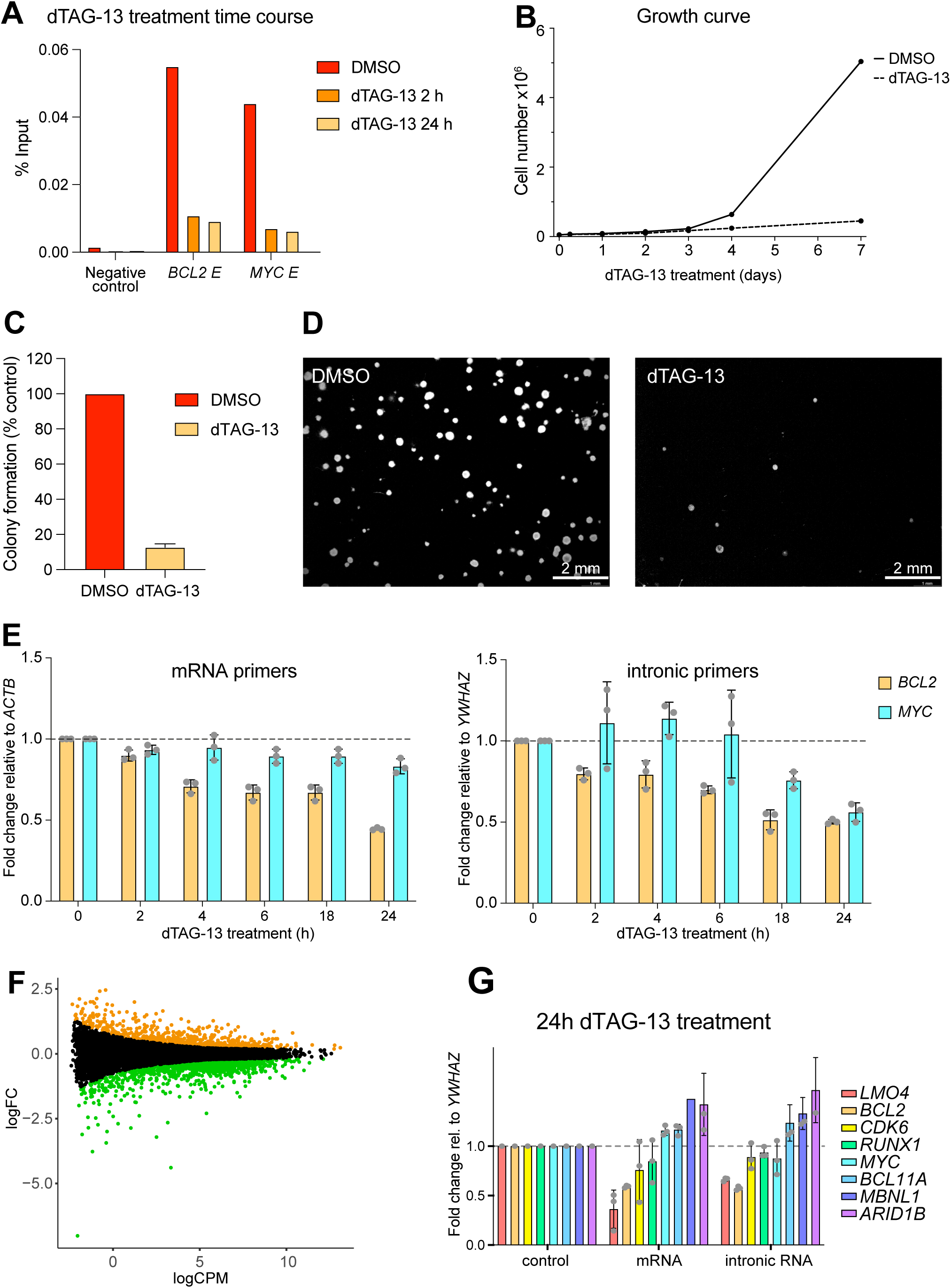
(**A**) ChIP-qPCR for MYB in untreated (DMSO) MYB-FKBP12^F36V^ tagged SEM cells or following treatment with 0.5 μM dTAG-13 for 2 h or 24 h. Representative of three biological replicates. (**B**) Growth of untreated (DMSO) or 0.5 μM dTAG-13 treated MYB-FKBP12^F36V^ tagged SEM cells over time. Representative of three biological replicates. (**C**) Colony counts 14 days after plating 500 cells per condition (DMSO vs treated with 0.5 μM dTAG-13 for 24 h) onto 1 ml semisolid methylcellulose medium. Data are the mean ± SD of three independent experiments. Three replicates were plated per experiment. (**D**) Representative images of colonies quantified in (C). (**E**) qRT-PCR analysis of *BCL2* and *MYC* expression following 0.5 μM dTAG-13 treatment for the indicated times, using primers to amplify mature mRNA (left) or intronic RNA (right). Values are normalized to *ACTB* (mature mRNA) or *YWHAZ* (intronic RNA), relative to DMSO treatment. Mean of three biological replicates; error bars show SEM. (**F**) Changes in TT-seq levels following 24h treatment with dTAG-13. Statistically significant differences (green: decreased; orange: increased; black: unchanged) from three biological replicates, FDR < 0.05. (**G**) qRT-PCR analysis of gene expression using mature mRNA and intronic PCR primers following 24h treatment with dTAG-13. Values are normalized *toYWHAZ* mature mRNA levels, relative to DMSO treatment. Mean of three biological replicates; error bars show SEM.

**Fig. S3.**
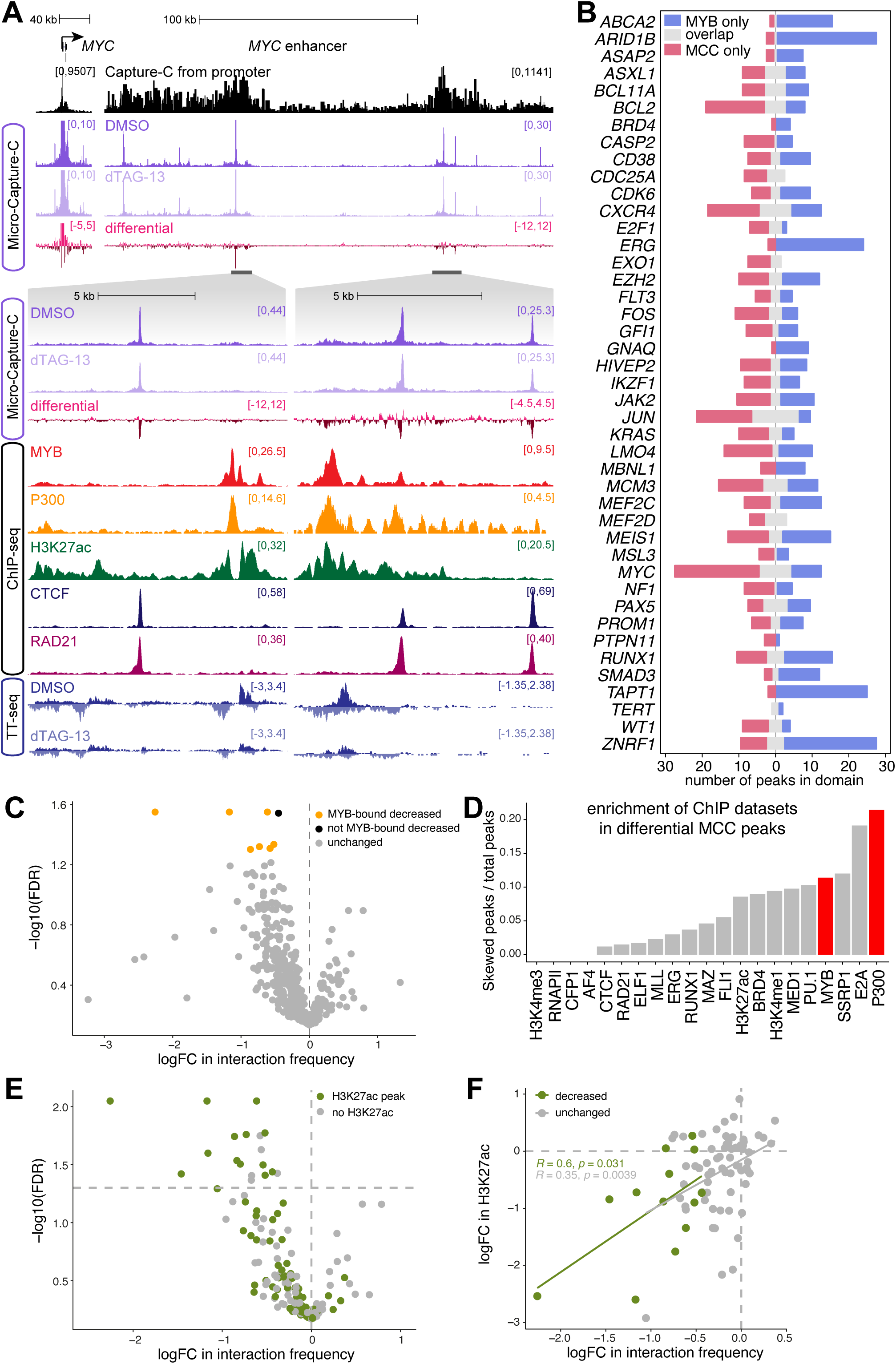
(**A**) Capture-C and ChIP-seq in SEM cells, and Micro-Capture-C (MCC) and TT-seq in MYB-FKBP12^F36V^ tagged SEM cells, with or without the addition of dTAG-13 for 24 h at the *MYC* locus. Capture-C and MCC traces scaled to emphasize distal interactions. (**B**) Overlap of MCC promoter-interacting peaks and MYB peaks within ∼500 kb of each gene indicated. (**C**) Change in interaction frequency at all MCC peaks showing interaction with target promoters following 24 h treatment with dTAG-13 in MYB-FKBP12^F36V^ tagged SEM cells. Statistically significant differences (yellow: decreased MYB bound peaks; black: decreased non-MYB-bound peaks; gray: unchanged) from three biological replicates, FDR <0.05. (**D**) Enrichment of SEM ChIP-seq datasets within differential MCC peaks. Statistically significant (FDR <0.05) MCC peaks with ChIP factor bound divided by total number of MCC peaks analysed with ChIP factor bound. MYB and its interaction partner P300 are highlighted in red. (**E**) Change in interaction frequency at all MCC peaks bound by MYB, following 24 h treatment with dTAG-13 in MYB-FKBP12^F36V^ tagged SEM cells. Peaks enriched for H3K27ac are shown in green. (**F**) Correlation of the change in promoter-interaction frequency and the change in H3K27ac at MYB-bound MCC peaks, following 24 h treatment with dTAG-13 in MYB-FKBP12^F36V^ tagged SEM cells. Spearman correlation coefficients are shown for peaks that show a statistically significant decrease in interaction frequency (FDR <0.05, green) or where interaction frequency is unchanged (gray).

**Fig. S4.**
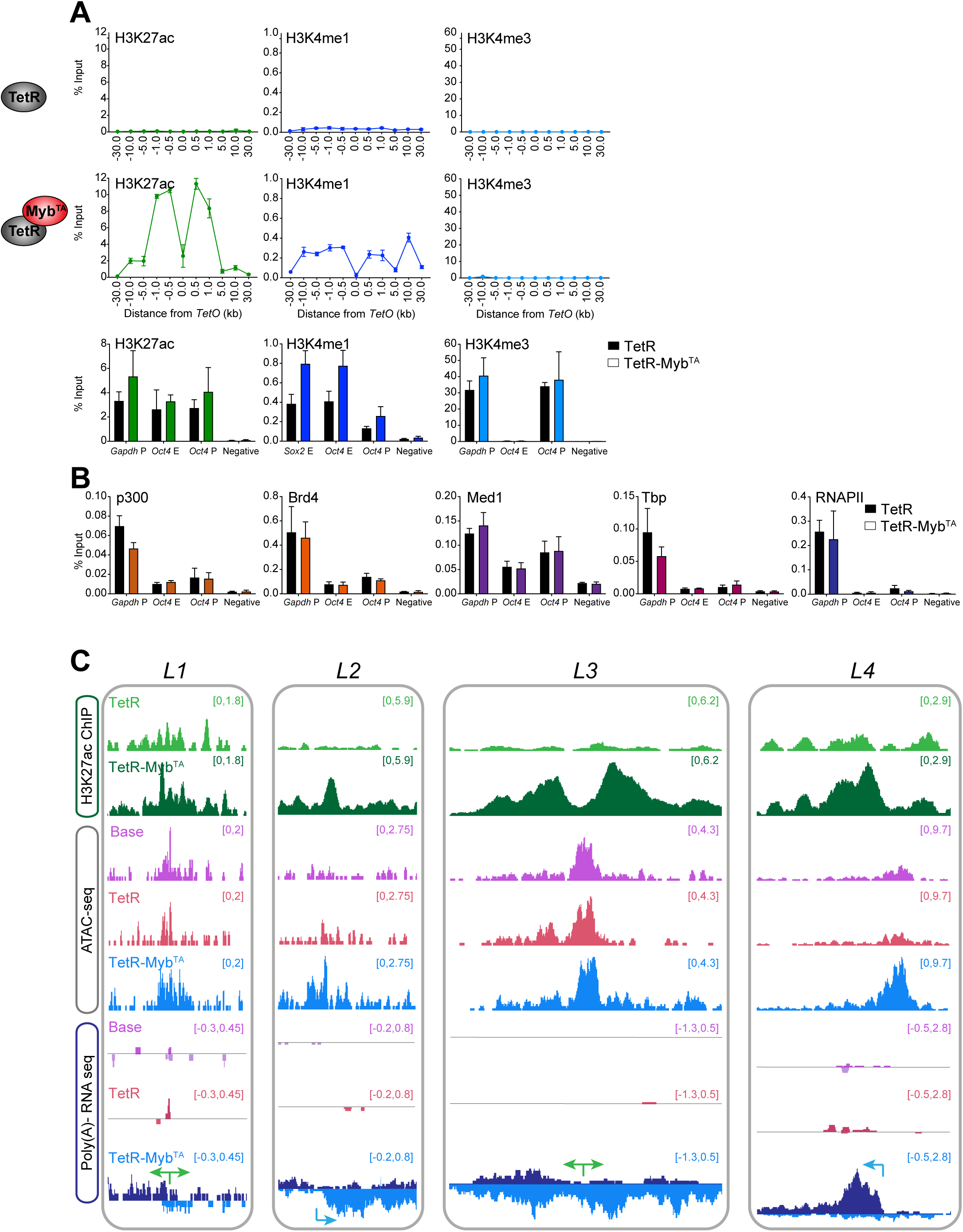
(**A**) ChIP-qPCR analysis across the *TetO*-containing locus in cell lines stably expressing either TetR (top row) or TetR-Myb^TA^ (middle row) and at endogenous loci (bottom row), for H3K27ac (left), H3K4me1 (middle) and H3K4me3 (right). Error bars represent the standard deviation from three biological replicates. (**B**) ChIP-qPCR for p300, Brd4, Med1, Tbp, RNAPII at endogenous loci in the indicated TetR mESCs. Error bars represent the standard deviation from three biological replicates. (**C**) H3K27ac ChIP-seq, ATAC-seq and poly-A minus RNA-seq at *L1-4* loci. Arrows represent predominant direction of transcription (green = bidirectional; blue = unidirectional).

**Fig. S5.**
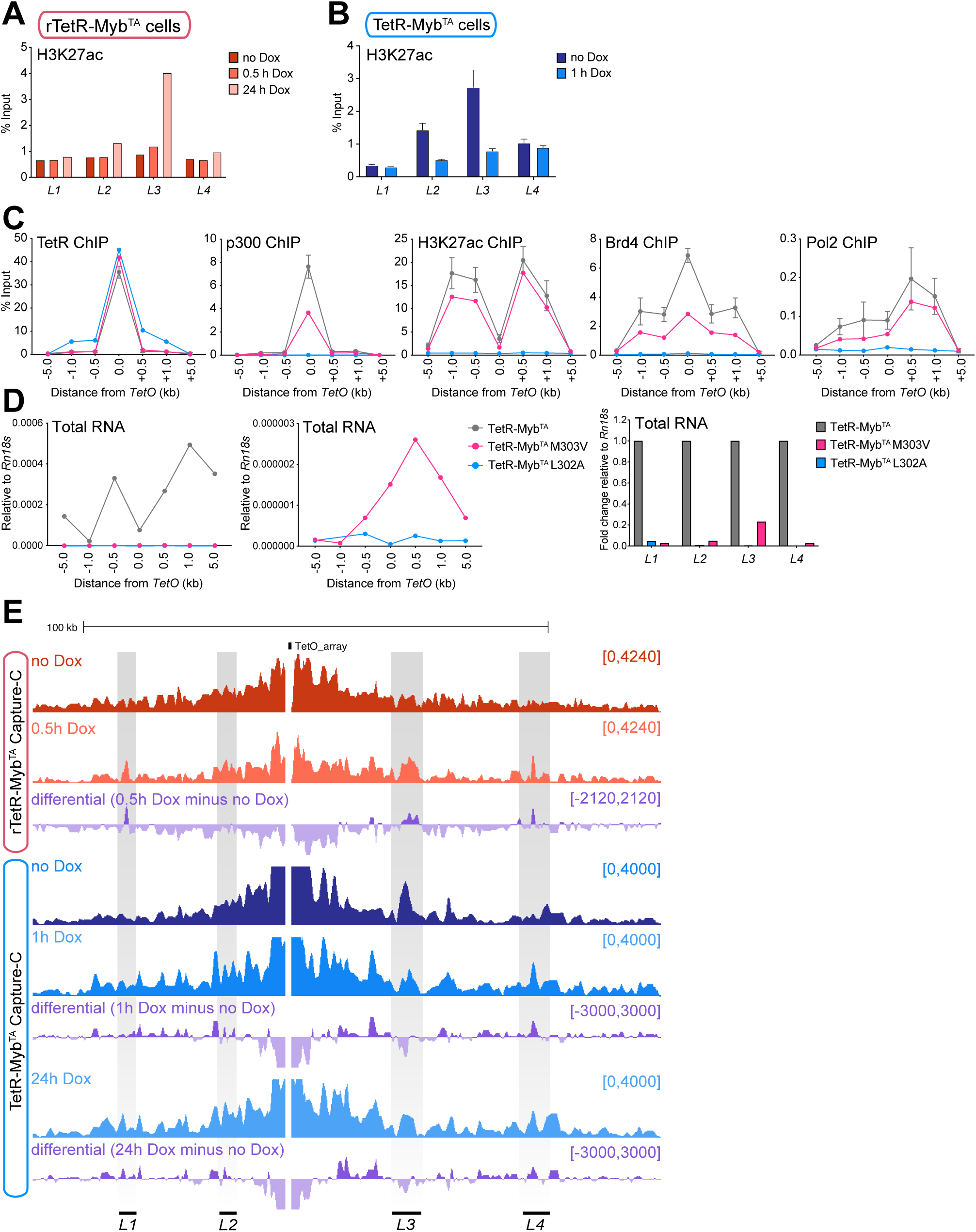
(**A**) ChIP-qPCR for H3K27ac at *L1-4* following doxycycline treatment for 0.5 and 24 h in rTetR-Myb^TA^ expressing cells. Representative of three biological replicates. (**B**) As in (A), ChIP-qPCR in TetR-Myb^TA^ expressing cells following doxycycline treatment for 1 h. Error bars represent the standard deviation from three biological replicates. (**C**) ChIP-qPCR analysis across the *TetO* containing locus in cell lines stably expressing either TetR-MYB^TA^ (gray), TetR-Myb^TA^ M303V (pink), or TetR-Myb^TA^ L302A (blue), using antibodies against TetR, p300, H3K27ac, Brd4 and RNAPII. Error bars represent the standard deviation from three biological replicates. (**D**) qRT-PCR across the *TetO* containing locus and at *L1-4* in TetR-MYB^TA^ (gray), TetR-Myb^TA^ M303V (pink), or TetR-Myb^TA^ L302A (blue) expressing cells. Representative of three biological replicates. (**E**) Capture-C across the *TetO* locus in rTetR-Myb^TA^ and TetR-Myb^TA^ expressing cells following treatment with doxycycline. Differential track shows the difference in interaction frequency between untreated and doxycycline conditions.

**Supplementary Table S1.**
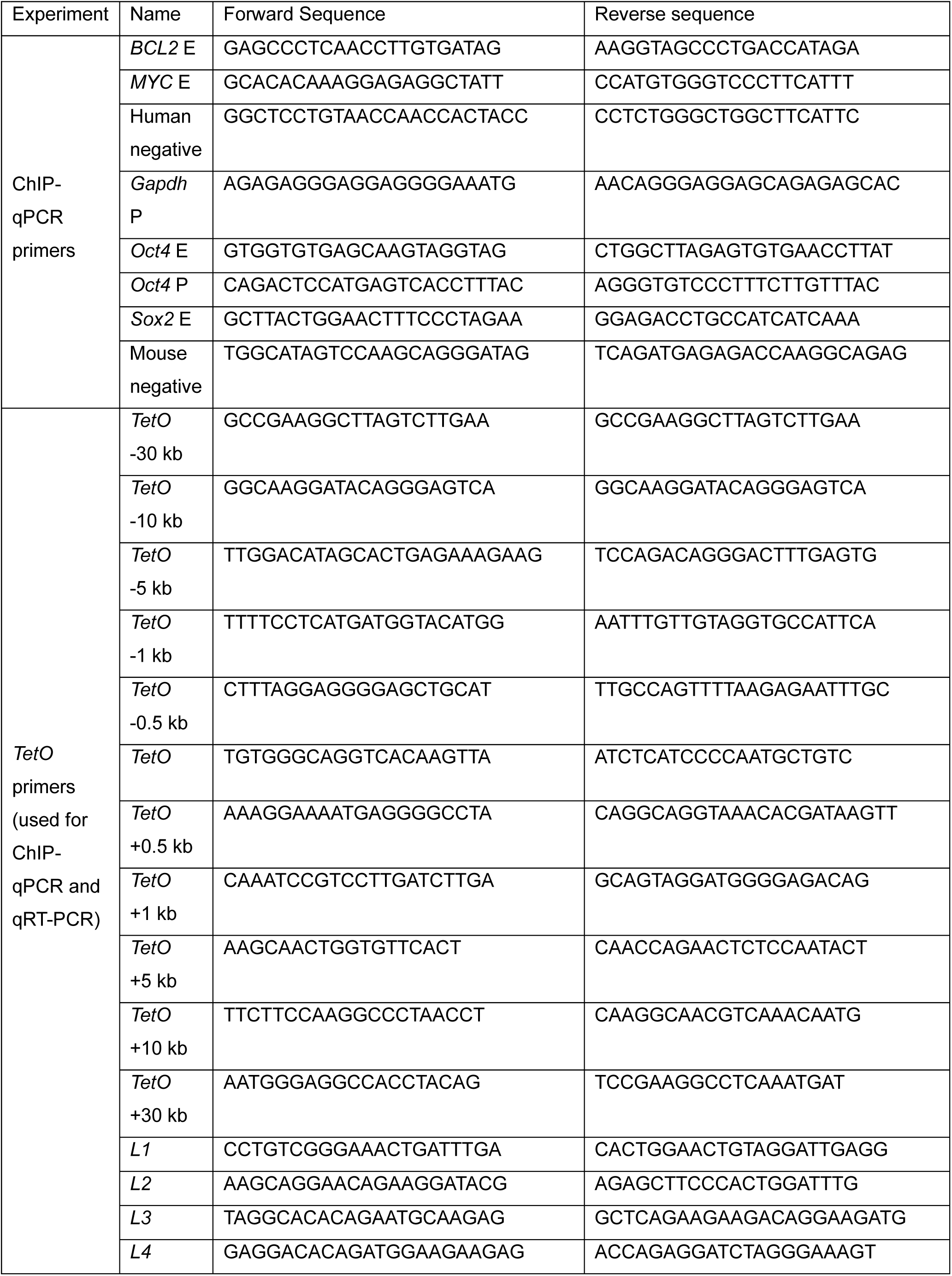
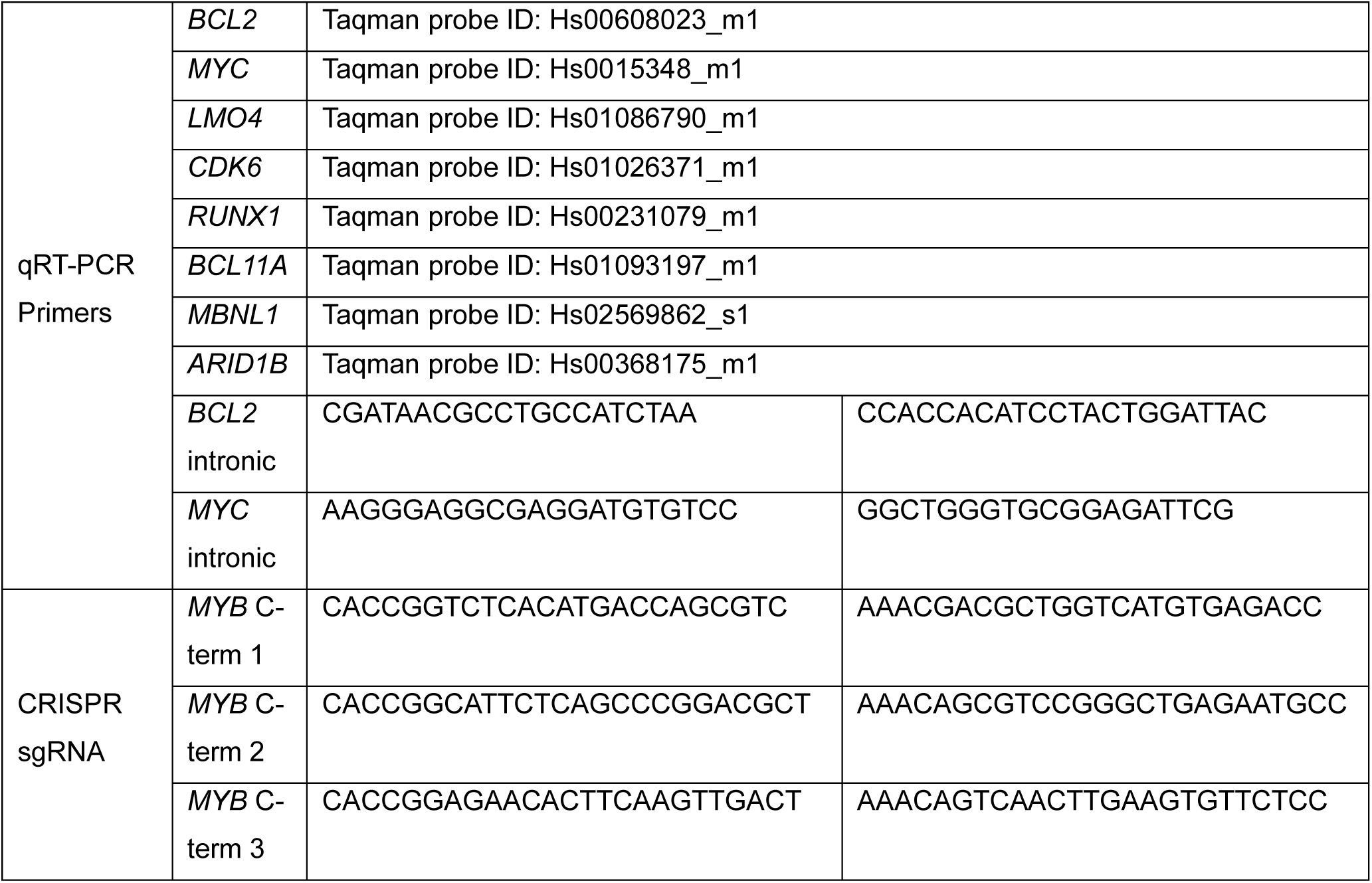
Primers and oligonucleotides used in this study.

**Supplementary Table S2.**
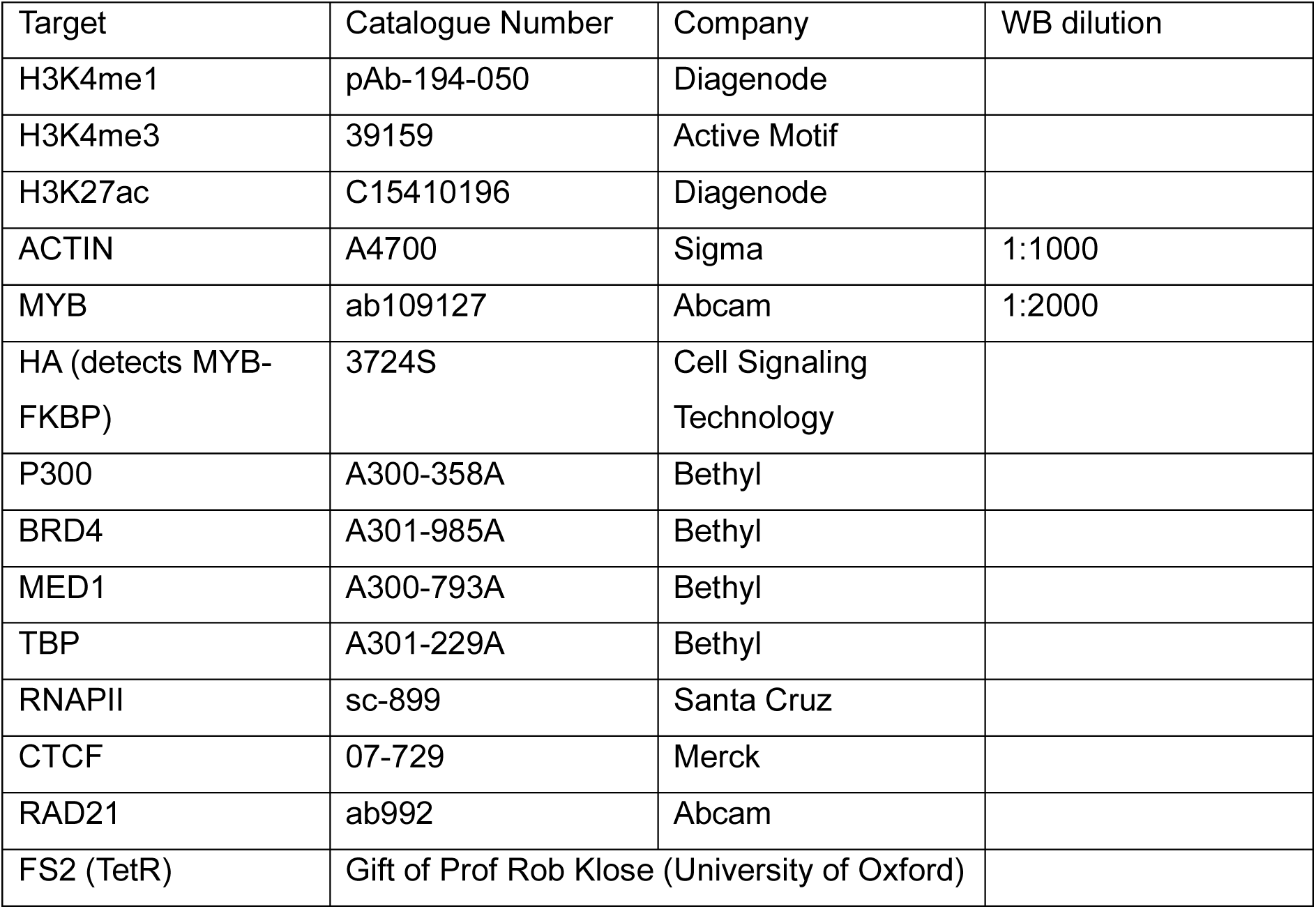
Antibodies used in this study.

| Target | Catalogue Number | Company | WB dilution |
| --- | --- | --- | --- |
| H3K4me1 | pAb-194-050 | Diagenode |  |
| H3K4me3 | 39159 | Active Motif |  |
| H3K27ac | C15410196 | Diagenode |  |
| ACTIN | A4700 | Sigma | 1:1000 |
| MYB | ab109127 | Abcam | 1:2000 |
| HA (detects MYB-FKBP) | 3724S | Cell Signaling Technology |  |
| P300 | A300-358A | Bethyl |  |
| BRD4 | A301-985A | Bethyl |  |
| MED1 | A300-793A | Bethyl |  |
| TBP | A301-229A | Bethyl |  |
| RNAPII | sc-899 | Santa Cruz |  |
| CTCF | 07-729 | Merck |  |
| RAD21 | ab992 | Abcam |  |
| FS2 (TetR) | Gift of Prof Rob Klose (University of Oxford) |  |  |

**Supplementary Table S3.**
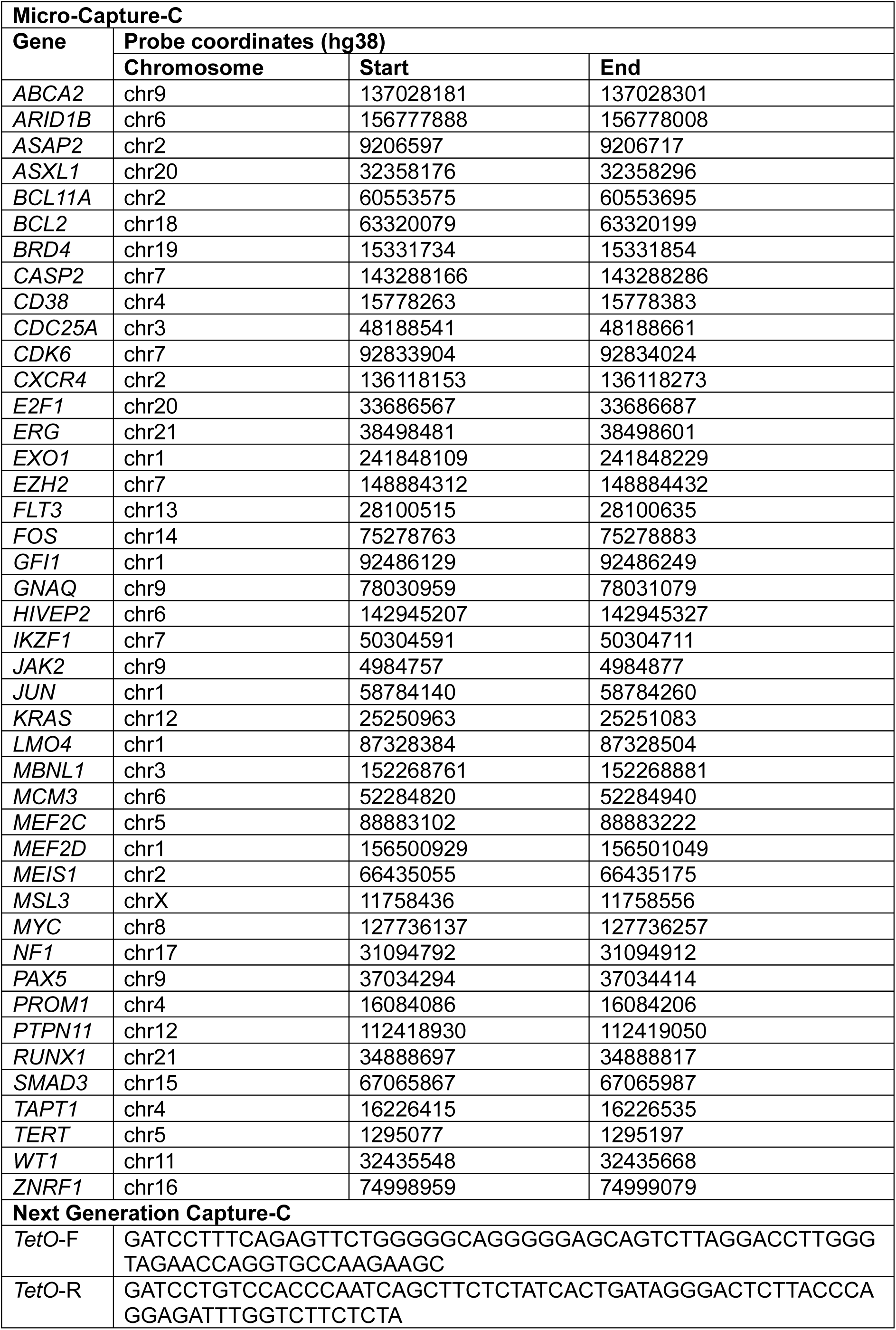
Probes used for NG Capture-C and Micro-Capture-C.

## References

1. F. Spitz, E. E. M. Furlong, Transcription factors: from enhancer binding to developmental control. Nature Reviews Genetics 13, 613--626 (2012).

2. Warren A. Whyte et al., Master Transcription Factors and Mediator Establish Super-Enhancers at Key Cell Identity Genes. Cell 153, 307--319 (2013).

3. Hannah K. Long, Sara L. Prescott, J. Wysocka, Ever-Changing Landscapes: Transcriptional Enhancers in Development and Evolution. Cell 167, 1170--1187 (2016).

4. M. Kassouf, S. Ford, J. Blayney, D. Higgs, Understanding fundamental principles of enhancer biology at a model locus: Analysing the structure and function of an enhancer cluster at the alpha-globin locus. Bioessays 45, e2300047 (2023).

5. J. H. Yang, A. S. Hansen, Enhancer selectivity in space and time: from enhancer-promoter interactions to promoter activation. Nat Rev Mol Cell Biol, (2024).

6. N. D. Heintzman et al., Distinct and predictive chromatin signatures of transcriptional promoters and enhancers in the human genome. Nat Genet 39, 311–318 (2007).

7. T. Zhang, Z. Zhang, Q. Dong, J. Xiong, B. Zhu, Histone H3K27 acetylation is dispensable for enhancer activity in mouse embryonic stem cells. Genome Biol 21, 45 (2020).

8. M. P. Creyghton et al., Histone H3K27ac separates active from poised enhancers and predicts developmental state. Proc Natl Acad Sci U S A 107, 21931–21936 (2010).

9. J. M. Alexander et al., Live-cell imaging reveals enhancer-dependent Sox2 transcription in the absence of enhancer proximity. Elife 8, (2019).

10. N. S. Benabdallah et al., Decreased Enhancer-Promoter Proximity Accompanying Enhancer Activation. Mol Cell 76, 473–484 e477 (2019).

11. A. Sanyal, B. R. Lajoie, G. Jain, J. Dekker, The long-range interaction landscape of gene promoters. Nature 489, 109–113 (2012).

12. E. E. M. Furlong, M. Levine, Developmental enhancers and chromosome topology. Science 361, 1341--1345 (2018).

13. J. Hyle et al., Acute depletion of CTCF directly affects MYC regulation through loss of enhancer-promoter looping. Nucleic Acids Res 47, 6699–6713 (2019).

14. L. F. Chen, J. Lee, A. Boettiger, Recent progress and challenges in single-cell imaging of enhancer-promoter interaction. Curr Opin Genet Dev 79, 102023 (2023).

15. Z. Chen et al., Increased enhancer-promoter interactions during developmental enhancer activation in mammals. Nat Genet 56, 675–685 (2024).

16. T. Pollex et al., Chromatin gene-gene loops support the cross-regulation of genes with related function. Mol Cell 84, 822–838.e828 (2024).

17. N. G. Aboreden et al., LDB1 establishes multi-enhancer networks to regulate gene expression. Mol Cell 85, 376–393 e379 (2025).

18. D. Hnisz et al., Super-enhancers in the control of cell identity and disease. Cell 155, 934–947 (2013).

19. J. Loven et al., Selective inhibition of tumor oncogenes by disruption of super-enhancers. Cell 153, 320–334 (2013).

20. N. T. Crump et al., BET inhibition disrupts transcription but retains enhancer-promoter contact. Nature Communications 12, 223 (2021).

21. L. El Khattabi et al., A Pliable Mediator Acts as a Functional Rather Than an Architectural Bridge between Promoters and Enhancers. Cell 178, 1145–1158 e1120 (2019).

22. S. Ramasamy et al., The Mediator complex regulates enhancer-promoter interactions. Nature Structural \& Molecular Biology 30, 991--1000 (2023).

23. P. Hua et al., Defining genome architecture at base-pair resolution. Nature 595, 125--129 (2021).

24. M. Maresca et al., Pioneer activity distinguishes activating from non-activating SOX2 binding sites. The EMBO Journal, e113150 (2023).

25. V. Y. Goel, M. K. Huseyin, A. S. Hansen, Region Capture Micro-C reveals coalescence of enhancers and promoters into nested microcompartments. Nat Genet 55, 1048–1056 (2023).

26. A. Agarwal, S. Korsak, A. Choudhury, D. Plewczynski, The dynamic role of cohesin in maintaining human genome architecture. Bioessays 45, e2200240 (2023).

27. M. L. Mucenski et al., A functional c-myb gene is required for normal murine fetal hepatic hematopoiesis. Cell 65, 677--689 (1991).

28. R. Sumner, A. Crawford, M. Mucenski, J. Frampton, Initiation of adult myelopoiesis can occur in the absence of c-Myb whereas subsequent development is strictly dependent on the transcription factor. Oncogene 19, 3335--3342 (2000).

29. H. Sakamoto et al., Proper levels of c-Myb are discretely defined at distinct steps of hematopoietic cell development. Blood 108, 896--903 (2006).

30. M. L. Sandberg et al., c-Myb and p300 Regulate Hematopoietic Stem Cell Proliferation and Differentiation. Developmental Cell 8, 153--166 (2005).

31. P. Dai et al., CBP as a transcriptional coactivator of c-Myb. Genes & Development 10, 528--540 (1996).

32. D. R. Pattabiraman, J. Sun, D. H. Dowhan, S. Ishii, T. J. Gonda, Mutations in Multiple Domains of c-Myb Disrupt Interaction with CBP/p300 and Abrogate Myeloid Transforming Ability. Molecular Cancer Research 7, 1477--1486 (2009).

33. P. Papathanasiou et al., A recessive screen for genes regulating hematopoietic stem cells. Blood 116, 5849--5858 (2010).

34. J. Zuber et al., An integrated approach to dissecting oncogene addiction implicates a Myb-coordinated self-renewal program as essential for leukemia maintenance. Genes \& Development 25, 1628--1640 (2011).

35. B. Biersack, M. Höpfner, Emerging role of MYB transcription factors in cancer drug resistance. Cancer Drug Resist 7, 15 (2024).

36. M. R. Mansour et al., An oncogenic super-enhancer formed through somatic mutation of a noncoding intergenic element. Science 346, 1373--1377 (2014).

37. J. O. J. Davies et al., Multiplexed analysis of chromosome conformation at vastly improved sensitivity. Nature Methods 13, 74--80 (2016).

38. B. Nabet et al., The dTAG system for immediate and target-specific protein degradation. Nat Chem Biol 14, 431–441 (2018).

39. B. Schwalb et al., TT-seq maps the human transient transcriptome. Science 352, 1225--1228 (2016).

40. J. M. Benito et al., MLL-Rearranged Acute Lymphoblastic Leukemias Activate BCL-2 through H3K79 Methylation and Are Sensitive to the BCL-2-Specific Antagonist ABT-199. Cell Rep 13, 2715–2727 (2015).

41. N. T. Crump et al., MLL-AF4 cooperates with PAF1 and FACT to drive high-density enhancer interactions in leukemia. Nat Commun 14, 5208 (2023).

42. N. P. Blackledge et al., Variant PRC1 complex-dependent H2A ubiquitylation drives PRC2 recruitment and polycomb domain formation. Cell 157, 1445–1459 (2014).

43. J. Kerry et al., MLL-AF4 Spreading Identifies Binding Sites that Are Distinct from Super-Enhancers and that Govern Sensitivity to DOT1L Inhibition in Leukemia. Cell Reports 18, 482--495 (2017).

44. H. Sakura et al., Delineation of three functional domains of the transcriptional activator encoded by the c-myb protooncogene. Proceedings of the National Academy of Sciences 86, 5758--5762 (1989).

45. K. Weston, J. M. Bishop, Transcriptional activation by the v-myb oncogene and its cellular progenitor, c-myb. Cell 58, 85--93 (1989).

46. F. Kalkbrenner, S. Guehmann, K. Moelling, Transcriptional activation by human c-myb and v-myb genes. Oncogene 5, 657–661 (1990).

47. J. Ernst et al., Mapping and analysis of chromatin state dynamics in nine human cell types. Nature 473, 43–49 (2011).

48. M. Gossen et al., Transcriptional Activation by Tetracyclines in Mammalian Cells. Science 268, 1766--1769 (1995).

49. D. Parker et al., Role of Secondary Structure in Discrimination between Constitutive and Inducible Activators. Molecular and Cellular Biology 19, 5601--5607 (1999).

50. T. Zor, R. N. D. Guzman, H. J. Dyson, P. E. Wright, Solution Structure of the KIX Domain of CBP Bound to the Transactivation Domain of c-Myb. Journal of Molecular Biology 337, 521--534 (2004).

51. E. Georgiades et al., Active regulatory elements recruit cohesin to establish cell-specific chromatin domains. bioRxiv, 2023.2010.2013.562171 (2023).

52. M. Gossen et al., Transcriptional activation by tetracyclines in mammalian cells. Science 268, 1766–1769 (1995).

53. L. Godfrey et al., DOT1L inhibition reveals a distinct subset of enhancers dependent on H3K79 methylation. Nature Communications 10, 2803 (2019).

54. J. D. Buenrostro, P. G. Giresi, L. C. Zaba, H. Y. Chang, W. J. Greenleaf, Transposition of native chromatin for fast and sensitive epigenomic profiling of open chromatin, DNA-binding proteins and nucleosome position. Nature Methods 10, 1213--1218 (2013).

55. B. Langmead, C. Trapnell, M. Pop, S. L. Salzberg, Ultrafast and memory-efficient alignment of short DNA sequences to the human genome. Genome Biol 10, R25 (2009).

56. H. Li et al., The Sequence Alignment/Map format and SAMtools. Bioinformatics 25, 2078–2079 (2009).

57. S. Heinz et al., Simple Combinations of Lineage-Determining Transcription Factors Prime cis-Regulatory Elements Required for Macrophage and B Cell Identities. Molecular Cell 38, 576--589 (2010).

58. W. J. Kent et al., The human genome browser at UCSC. Genome Res 12, 996–1006 (2002).

59. F. Ramírez et al., deepTools2: a next generation web server for deep-sequencing data analysis. Nucleic Acids Research 44, W160--W165 (2016).

60. D. Kim, J. M. Paggi, C. Park, C. Bennett, S. L. Salzberg, Graph-based genome alignment and genotyping with HISAT2 and HISAT-genotype. Nat Biotechnol 37, 907–915 (2019).

61. Y. Liao, G. K. Smyth, W. Shi, featureCounts: an efficient general purpose program for assigning sequence reads to genomic features. Bioinformatics 30, 923--930 (2014).

62. Y. Chen, L. Chen, A. T. L. Lun, P. L. Baldoni, G. K. Smyth, edgeR v4: powerful differential analysis of sequencing data with expanded functionality and improved support for small counts and larger datasets. Nucleic Acids Res 53, (2025).

63. D. J. Downes et al., Capture-C: a modular and flexible approach for high-resolution chromosome conformation capture. Nat Protoc 17, 445–475 (2022).

64. J. C. Hamley, H. Li, N. Denny, D. Downes, J. O. J. Davies, Determining chromatin architecture with Micro Capture-C. Nat Protoc 18, 1687–1711 (2023).

65. L. D. Hentges et al., LanceOtron: a deep learning peak caller for genome sequencing experiments. Bioinformatics 38, 4255–4263 (2022).

